# Evolved, Selective Erasers of Distinct Lysine Acylations

**DOI:** 10.1101/723684

**Authors:** Martin Spinck, Maria Ecke, Raphael Gasper, Heinz Neumann

## Abstract

Lysine acetylation, including related lysine modifications such as butyrylation and crotonylation, is a widespread post-translational modification with important roles in many important physiological processes. However, uncovering the regulatory mechanisms that govern the reverse process, deacylation, has been challenging to address, in great part because the small set of lysine deacetylases (KDACs) that remove the modifications are promiscuous in their substrate and acylation-type preference. This lack of selectivity hinders a broader understanding of how deacylation is regulated at the cellular level and how it is correlated with lysine deacylation-related diseases. To facilitate the dissection of KDACs with respect to substrate specificity and modification type, it would be beneficial to re-engineer KDACs to be selective towards a given substrate and/or modification. To dissect the differential contributions of various acylations to cell physiology, we developed a novel directed evolution approach to create selective KDAC variants that are up to 400-fold selective towards butyryl- over crotonyl-lysine substrates. Structural analyses of this non-promiscuous KDAC revealed unprecedented insights regarding the conformational changes mediating the gain in specificity. As a second case study to illustrate the power of this approach, we re-engineer the human SirT1 to increase its selectivity towards acetylated versus crotonylated substrates. These new enzymes, as well as the generic approach that we report here, will greatly facilitate the dissection of the differential roles of lysine acylation in cell physiology.

**Significance Statement:** Acetylation of lysine residues features numerous roles in diverse physiological processes and correlates with the manifestation of metabolic diseases, cancer and ageing. The already huge diversity of the acetylome is multiplied by variations in the types of acylation. This complexity is in stark contrast to the small set of lysine deacetylases (KDACs) present in human cells, anticipating a pronounced substrate promiscuity.

We device a strategy to tackle this disarray by creating KDAC variants with increased selectivity towards particular types of lysine acylations using a novel selection system. The variants facilitate the dissection of the differential contributions of particular acylations to gene expression, development and disease. Our structural analyses shed light on the mechanism of substrate discrimination by Sirtuin-type KDACs.

## Introduction

Protein acetylation was first discovered more than fifty years ago as a posttranslational modification of histone proteins (1). The past two decades revealed a wide variety of functional roles of this modification in almost every physiological process (2) such as transcription (3), chromatin structure (4), cytoskeleton organization (5) and energy metabolism (6). Unsurprisingly, defects in the enzymes governing lysine acetylation are linked to a variety of diseases such as diabetes (7), cancer (8), neurodegeneration (9), and aging (10). The acylation spectrum of lysine side chains is not restricted to acetylation, but broad, from short acyl chains to fatty acids and charged functional groups (11, 12). All of these modifications are reversed by a comparably small set of lysine deacetylases (KDACs) grouped into four enzyme families. The related class 1, 2 and 4 enzymes (13) are structurally and mechanistically distinct from class 3 KDACs (14, 15). The former classes contain an active site zinc ion to orient a water molecule and to polarize the substrate. The latter class, the Sirtuins, consist of a zinc binding and a Rossmann-fold domain, responsible for acyl peptide- and NAD^+^-binding, respectively (7), connected by a flexible cofactor binding loop (15, 16). They couple the hydrolysis of the amide bond to the cleavage of NAD^+^ (17), producing nicotinamide (NAM) and O-acetyl-ADP-ribose (18).

KDACs differ in their preference towards various acylations and peptide sequences, the biological relevance and purpose of which are still unclear (19). For example, class 1 KDACs (HDAC1, HDAC2, HDAC3 and HDAC8) and SirT1 exhibit significant decrotonylase activity *in vivo* (20). SirT1–3 are active towards various uncharged acylations *in vitro*, while SirT5 mainly removes negatively charged acylations and SirT6 and HDAC11 exhibit preference for long chain fatty acids (21–25).

Despite extensive structural and mechanistic studies (15, 26–28), how KDACs discriminate between different types of acylations remains poorly understood. In order to deconvolute the role of different lysine acylations, acyl-type selective KDAC variants would be tremendously useful tools. The attempt to create an acetyl-specific HDAC1 variant by rational design resulted in an enzyme that lost deacetylation and retained decrotonylation activity (19, 20). Hence, the outcome of attempts to design specificity in KDACs is difficult to predict.

Here, we report on the design of a selection system for KDACs enabling directed evolution of variants with altered deacylation selectivity. The system relies on incorporation of lysine derivatives by genetic code expansion (29) in the reporter enzyme, orotidine-5’-monophosphate (OMP) decarboxylase, in place of an essential lysine residue at the active site. The OMP decarboxylase containing the lysine derivative is an inactive precursor that turns on upon removal of the modification, thereby coupling deacetylase activity to a selectable output (Fig. 1A). This allows us to evolve KDACs selective for particular lysine acylations. These KDAC variants can be used to partially complement KDAC deletion strains to reveal the physiological role of alternative lysine acylations.

**Fig. 1.**
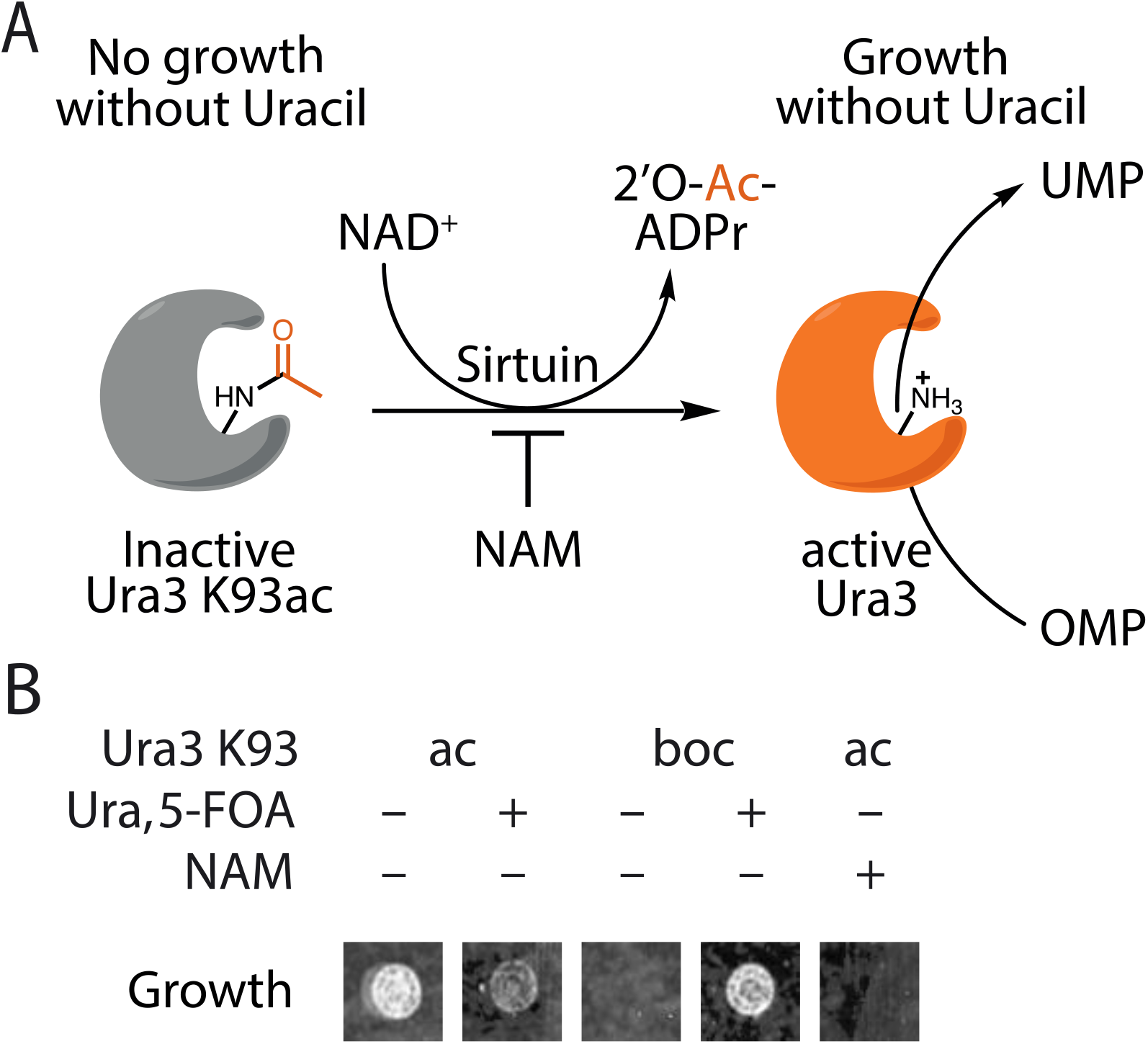
Design of the KDAC selection system. **A)** Deacetylation of OMP decarboxylase K93ac (Ura3 K93ac) by Sirtuins enables growth of *E. coli* in the absence of uracil. **B**) *E. coli* producing Ura3 K93ac as the sole source of OMP decarboxylase depend on KDAC activity. Cell growth of *E. coli* DB6656 (ΔpyrF) expressing plasmids to encode Ura3 K93ac or K93boc (not cleavable, tert-butyloxycarbonyl-lysine) on agar plates with or without uracil and 5-FOA. Nicotinamide (NAM), which inhibits endogenous CobB, prevents growth.

## Results

### Design of a selection system for acyl-type specific lysine deacetylases

In order to develop a selection system for acyl-type specific KDACs, we first searched for a selectable marker with an essential lysine residue that could be masked by acylation. Mutation of an active site lysine (K93) to alanine of budding yeast OMP decarboxylase (Ura3, which is required for the biosynthesis of uracil) is known to reduce its activity by more than five orders of magnitude (30). To test whether acetylation of K93 has a similar impact on catalysis, we complemented *E. coli* ΔpyrF (the homologue of Ura3) with Ura3 K93ac (Fig. 1B). Ura3 K93ac is produced by replacing the respective lysine codon in the Ura3 gene with an amber (UAG) codon and encoding the incorporation of N(ε)-acetyl-lysine (AcK) with *M. barkeri* AcKRS3/PylT (an AcK-specific variant of pyrrolysyl-tRNA synthetase (PylS) and its cognate amber suppressor tRNA (31, 32)).

We tested the selection system by cultivating *E. coli* expressing Ura3 K93ac as the sole source of OMP decarboxylase on minimal media without uracil. The cells were able to grow in the absence of uracil, but did not grow when CobB (the major lysine deacetylase of *E. coli*) was inhibited with NAM, indicating that CobB was able to remove the acetyl group from the active site lysine of Ura3 K93ac. Growth of the same cells is inhibited when 5-fluoro-orotic acid (5-FOA), a compound converted to a toxic metabolite by Ura3 (33), is added to the medium. Incorporation of N(ε)-tert.-butyl-oxycarbonyl-lysine (BocK) in place of AcK did not complement pyrF deficiency, indicating that this lysine modification is not a substrate of CobB. Hence, this system is able to positively and negatively select *E. coli* harboring an active KDAC. To demonstrate the portability of our selection system to other KDACs, we used it to select for human SirT1-3 and HDAC8 activity, a mammalian class I KDAC structurally and mechanistically distinct from the Sirtuin family member CobB (Supplementary Figure 1).

### Isolation of acyl-type specific deacetylases

Next, we aimed to create acyl-type selective variants of CobB. Therefore, we constructed a mutant library by randomizing five active site residues (A76, Y92, R95, I131 and V187) of CobB to all possible combinations of natural amino acids, thereby creating 20^5^ (3.2×10^6^) different mutants (Fig. 2A). To identify CobB mutants that selectively remove acetyl but not crotonyl groups, we subjected the library to three rounds of selection (positive, negative, positive). In the first round of positive selection for CobB variants able to remove acetyl groups, we grew *E. coli* ΔpyrF ΔcobB transformed with the CobB mutant library and expressing Ura3 K93ac in the presence of AcK on medium without uracil. The subsequent round of negative selection aimed to remove CobB variants active towards crotonyl groups from the remaining pool. Therefore, the library plasmids were isolated from the pool of surviving clones of the first round of selection and used to transform *E. coli* ΔpyrF ΔcobB producing Ura3 K93cr (encoded with wild-type PylS (34–36)). Cells were grown on plates containing N(ε)-crotonyl-lysine (CrK) and 5-FOA to select against CobB mutants capable of removing crotonyl groups from Ura3 K93cr. After a third positive round of selection for deacetylase activity, CobB library plasmids were isolated from individual clones and re-tested for their ability to allow cells to survive on uracil-free medium when expressing Ura3 K93ac.

**Fig. 2.**
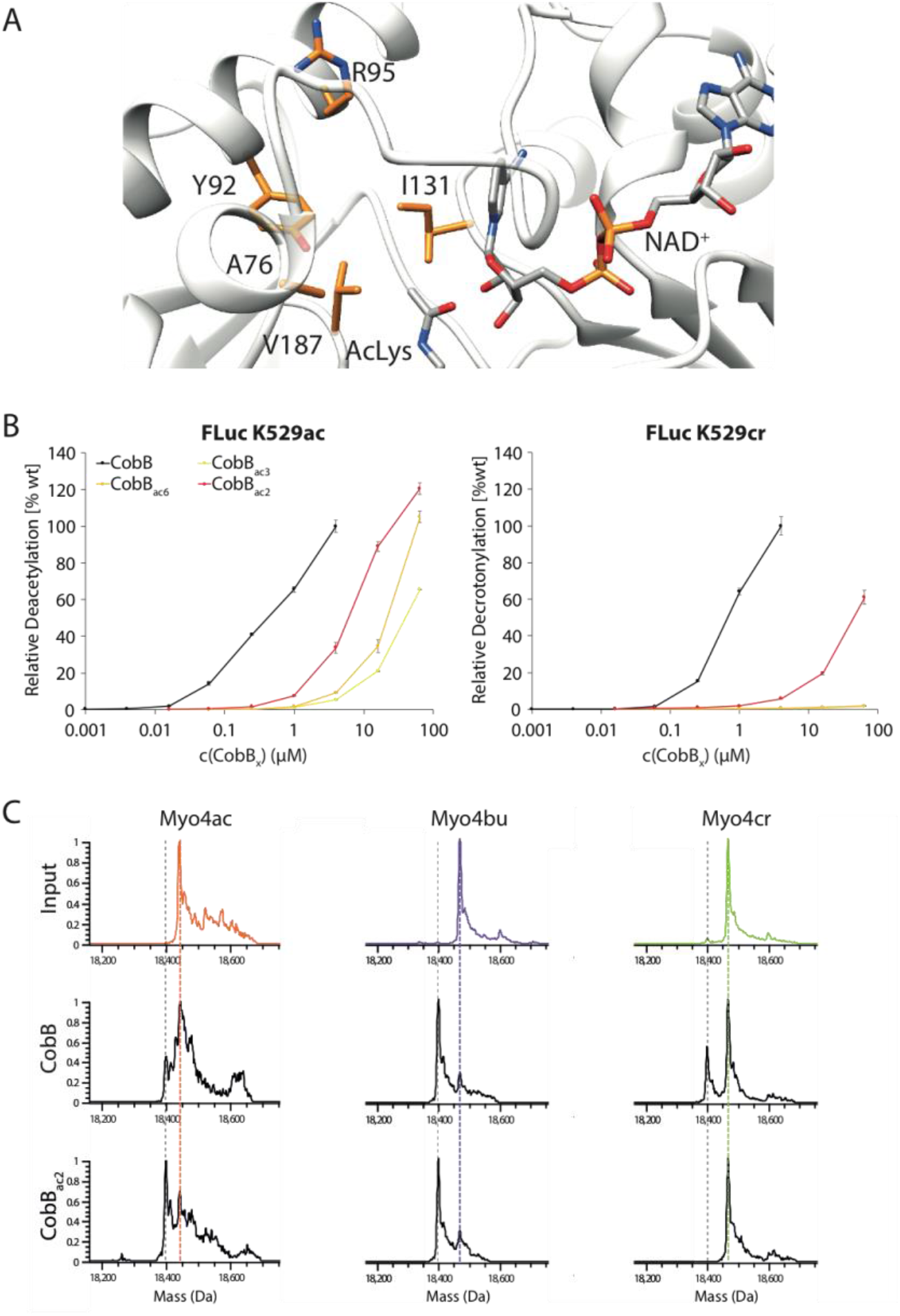
Creation of AcK-selective CobB variants. A) CobB library design. Highlighted amino acid residues were randomized to all possible combinations of natural amino acids to generate a library of thirty million mutants in the active site of CobB (image was created using pdb-file 1S5P). B) Demodification of FLuc K529ac and FLuc K529cr by CobB and AcK-selective variants. C) LC-ESI-MS analysis of myoglobin proteins before and after treatment with CobB or CobB_ac2_. Dotted lines mark the expected mass of the free (grey) or modified lysine (colored).

We sequenced 60 isolates corresponding to 14 unique CobB mutants. Eight of the sequences dominated this set and accounted for 53 of the sequenced CobB isolates (Table S1). Hence, the selection system is capable of enriching individual sequences from a mean frequency of 3.125×10^−7^ (1/3.2×10^6^) in the original library to 0.3 (18 out of 60) after three rounds of selection, suggesting amplification by a factor of almost one million.

To identify the most active KDAC variants among the isolates, we set out to modulate the stringency of our selection system. The reporter enzyme Ura3 is inhibited by 6-azauridine (6-AU). Addition of this compound to the medium should therefore raise the threshold level of this enzyme required for cell growth. We arrayed the 14 different clones isolated in the selection on plates containing increasing concentrations of 6-AU and monitored growth at 37°C (Supplementary Fig. 2). The ability of the clones to survive increasing 6-AU concentrations varied between 0.16 to 2 mM.

### Biochemical characterization of evolved CobB variants

Next, we purified the mutant enzymes and analyzed their ability to reverse the modification of purified FLuc K529ac, K529bu and K529cr (37) (Fig. 2B and Supplementary Figure 3). With 25% activity against FLuc K529ac and almost 100% against FLuc K529bu compared to wild-type, CobB_ac2_ turned out to be the most active variant, approximately fourfold more active than CobB_ac6_ (Fig. 2B), consistent with providing the highest resistance to 6-AU (Supplementary Fig. 2).

The diminished activity of CobB_ac3_ and CobB_ac6_ for AcK may be explained by an increased K_M_ for NAD^+^ (Supplementary Fig. 4A). These mutants retain similar sensitivity to inhibition by NAM. CobB_ac6_ shows a decreasing deacylation activity with increasing acyl chain length (Supplementary Fig. 4B). Interestingly, CobB_ac2_ and CobB_ac3_ prefer the longer acyl modification of BuK more than tenfold over AcK, resulting in a remarkable selectivity of these variants for BuK over CrK.

CobB_ac2_ still showed measurable decrotonylation activity compared to CobB_ac3/6_, however, when we tested this mutant using myoglobin with different acylations at residue four (32) as substrates, it was able to remove acetylation and butyrylation but not crotonylation (Fig. 2C). The more selective variants, CobB_ac3_ and CobB_ac6_, were unable to remove the acetylation (or any other acylation tested) of this residue to a measurable extent (Supplementary Figure 5). We attribute this lack of activity to the difference in the substrates and the differential sensitivity of the assays to detect the unmodified protein.

Similarly, CobB_ac2_ was capable of removing H4 K16ac from isolated HeLa histones, while its ability to reverse histone crotonylation was strongly reduced compared to wild-type CobB (Supplementary Fig. 6). Hence, our combined selection and screening system is able to identify acyl-type specific variants in a large pool of KDAC mutants. Particularly CobB_ac2_ could be used to shift the lysine acylation pattern of the proteome towards crotonylation or to selectively deplete lysine butyrylation, and thus may help to uncover the specific role of these modifications in cell physiology.

### Crystal structure of butyryl-specific CobB

To obtain insight into the mechanism of substrate selectivity, we crystalized CobB and CobB_ac2_ in complex with histone H4 peptides acylated at lysine 16 with either acetyl, butyryl or crotonyl (see Table S2 for crystallographic data). The overall structure of CobB_ac2_ is very similar to wild-type CobB with a rmsd of less than 0.5 Å (Figure 3). The most pronounced changes in the CobB_ac2_ structures bound to different peptides concentrate on the substrate binding loop connecting Helix α1 and Helix α2, whereas the wild-type structures are little affected by the type of acylation of the bound peptide (Figure 3).

**Fig. 3:**
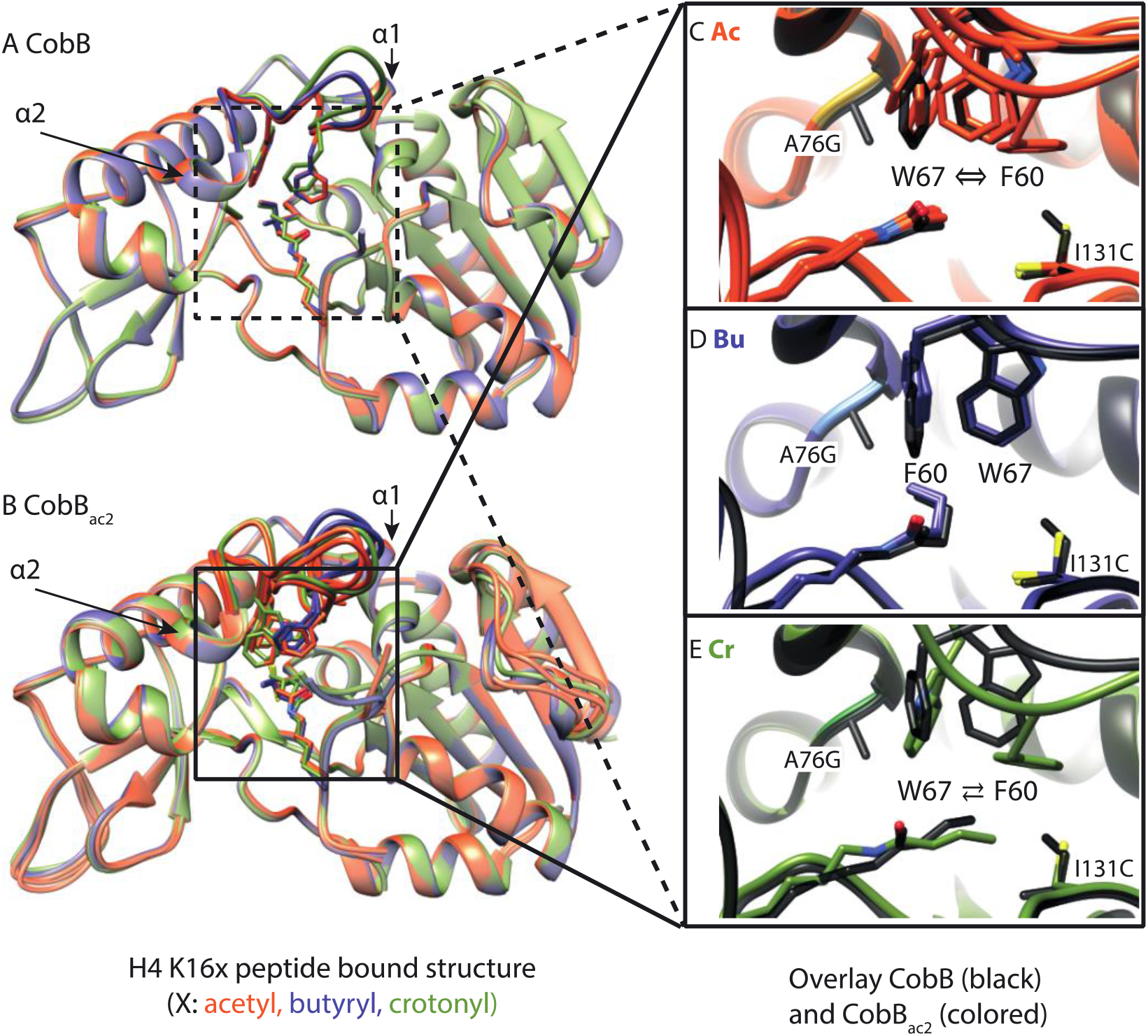
Crystal structures of wild-type CobB (A) and CobB_ac2_ (B, A76G, I131C) in complex with K16-modified H4 peptides (acetyl (red-brown), butyryl (blue), crotonyl (green)). Overlay of different substrates shows high conformational variability of the cofactor binding loop connecting Helix α1 and Helix α2 (highlighted) of CobB_ac2_ (B) compared to wild-type (A). C-E) Overlay of active site residues (wild-type black, CobBac2 colored) in the presence of the same substrate. Depending on the type of modification, the substrates either induce the native conformation (D, Bu), a flipped conformation (E, Cr, ⇄), or a mixture of both (C, Ac, ⇔) in CobB_ac2_. Several chains from an elementary cell are overlaid (for chains D and H in CobB_ac2_:H4 K16ac and chain A in CobB_ac2_:H4 K16cr see Supplementary Figure 7).

The mutations in CobB_ac2_, A76G and I131C, allow the loop to adopt two alternative conformations, the choice of which is dictated by the substrate. The loop conformations differ mainly in the arrangement of W67 and F60. The butyryl group stabilizes the original arrangement of these residues by its kinked binding mode. A crotonyl group lacks this flexibility due to its conjugated bonds and, therefore, dislocates F60 from its original position. The vacated space is then occupied by W67, causing a distortion of the cofactor binding loop (Figure 3E). Although the enzyme prefers to adopt this new conformation, it is still able to rearrange to its original conformation, because a second chain in the same unit cell shows a superposition of both structures (Supplementary Figure 7). The acetyl group does not stabilize either conformation, resulting in multiple arrangements of the loop observed for different chains within the same unit cell.

We observed similar structural alterations for CobB_ac3_ bound to the acetylated peptide (Supplementary Figure 8A, B), indicating that it acquires its selectivity by a similar mechanism. To test if swapping of F60 and W67 is the main cause of the butyryl selectivity we interchanged both residues by mutagenesis. CobB F60W W67F displayed a similar selectivity for butyryl lysine as CobB_ac2_ (Supplementary Figure 8C). Hence, this structural alteration is the key feature responsible for butyryl selectivity. However, as we could not obtain crystals of CobB_ac3_ bound to butyrylated or crotonylated peptides, we cannot exclude other explanations for the selectivity of CobB_ac3_.

Since we expected the altered conformation to be inactive, we attempted to solve the structure of the complex of CobB_ac2_ with NAD^+^ and crotonyl peptide (Figure 4). We obtained crystals of the complex by co-crystallization after 16 h (Figure 4A) and by soaking for 36 h (Figure 4B). We observed a continuous density that linked the ADP-ribose moiety to the crotonyl group via the 2’-OH of the ribose (Figure 4B), suggesting the presence of a reaction intermediate. This density is best explained by reaction intermediate III (Int. III) that was previously reported by Wang et al. for the SirT2 reaction cycle in the presence of a thiomyristoyl inhibitor (38). We modelled the analogous crotonyl-Int. III with 60% (A) and 100% (B) occupancy for both states. Usually, this intermediate is hydrolyzed, but the repositioned W67 blocks the access of water to the active site (Figure 4A). Indeed, water is only near Int. III after W67 has returned to its original position, as seen in the 36 h structure (Figure 4B). Nevertheless, there is no clear indication in this structure for hydrolysis of the intermediate, suggesting that other factors may also be required.

In summary, we have shown that CobB_ac2_ acquires its selectivity for butyrylated peptides through a substrate-induced mechanism in which the mutations A76G and I131C allow the cofactor binding loop to adopt two alternative conformations. The crotonyl induced conformation is inactive because access of water to the active site is blocked. This leads to an accumulation of Int. III, which is slowly hydrolyzed upon reversion of the loop conformation.

**Fig. 4.**
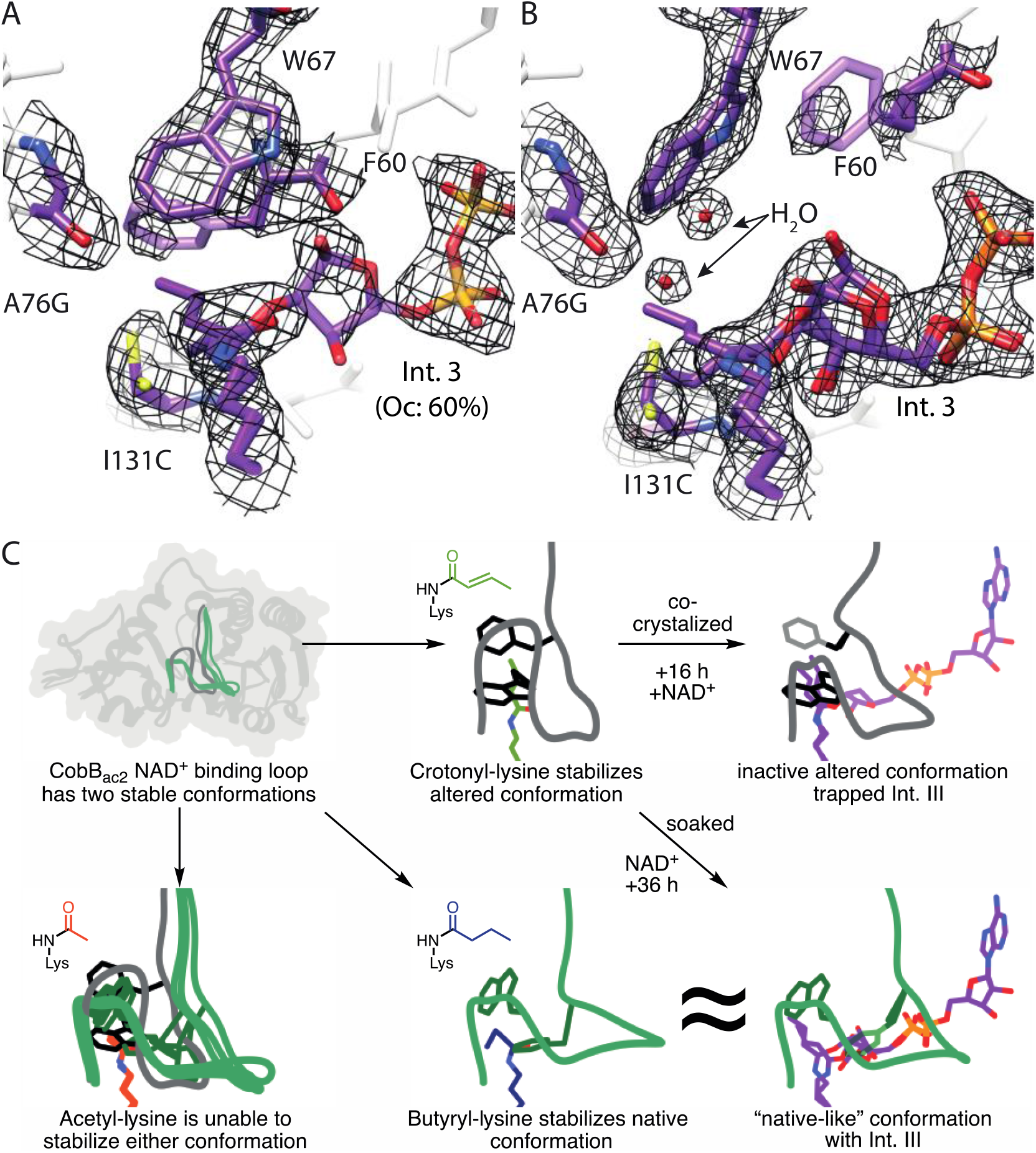
Demodification of crotonylated peptide stalls at intermediate 3 in the CobB_ac2_ mutant. Active site view of the crystal structure obtained after 16 h (A, crystalized CobB_ac2_, H4K16cr and NAD^+^) and 36 h (B, NAD^+^ soaked CobB_ac2_/H4 K16cr crystals). The 2Fo-Fc omit map (black, 1 σ) is shown for key residue. The density map shows a clear continuous density between the crotonyl group and the 2’-OH group of the ADP-ribose. Therefore, the intermediate was modelled analogously to the intermediate III described by Wang et al. 2017 (35). The cartoon (C) highlights the critical changes in the cofactor binding loop of CobB_ac2_ by which the enzyme recognizes and specifically stalls the decrotonylation reaction.

### Creation of acyl-type selective SirT1 variants

To test the generality of our selection approach, we created a library in the active site of SirT1 by randomizing five residues (A313, I316, I347, F366, I411) to all possible combinations of natural amino acids (Fig. 5A). We subjected this library to three rounds of selection (positive on Ura3 K93ac; negative on either Ura3 K93pr/bu/cr; positive on Ura3 K93ac) and sequenced in total 82 isolates after the third round (Fig. 5B).

**Fig. 5:**
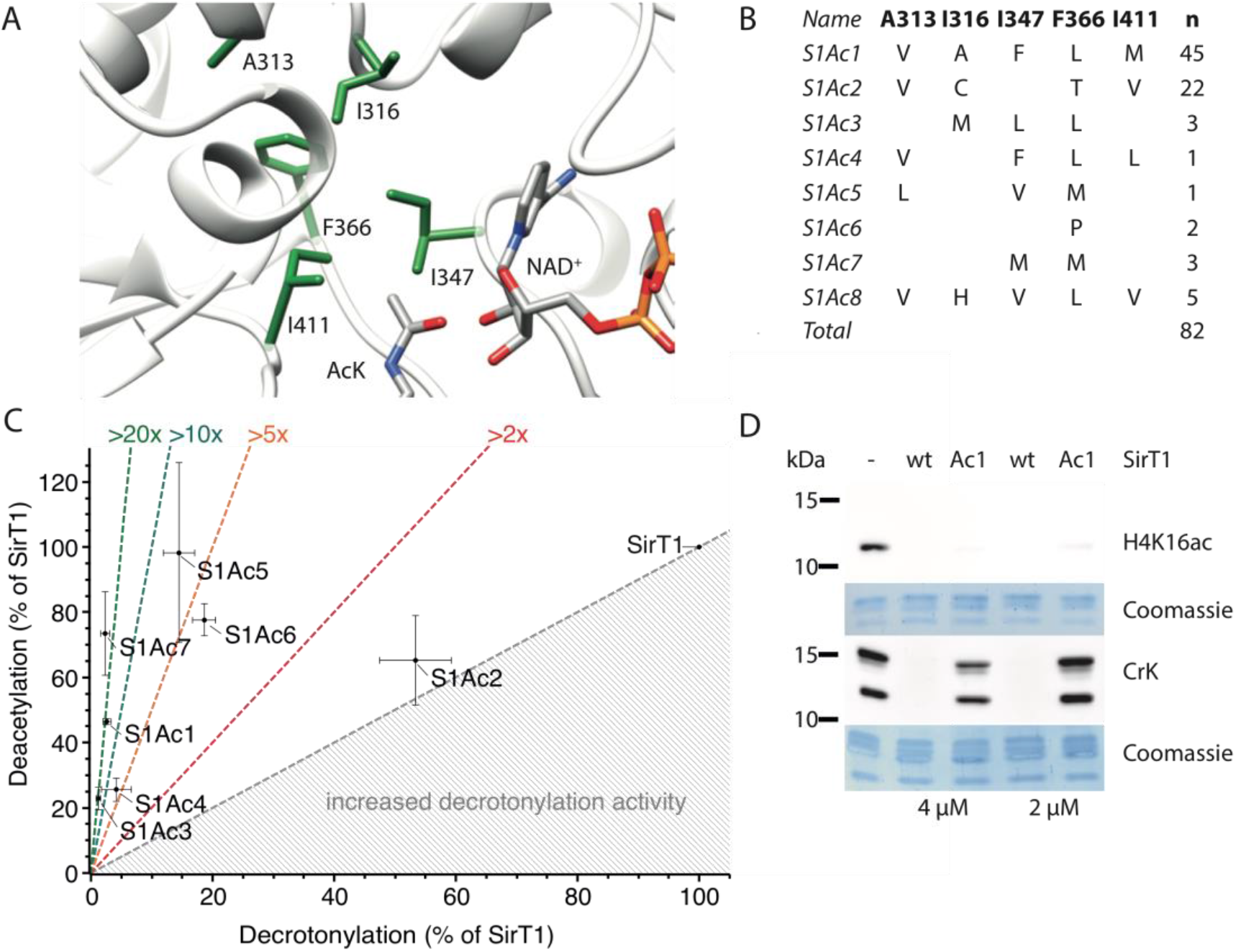
Creation of SirT1 variants with AcK preference. A) Structure of the SirT1 active site (4KXQ) with ligands NAD^+^ and AcK imported from pdb-files 2H4F and 1S5P, respectively, shown as sticks. Residues highlighted in green were randomized to all possible combinations of natural amino acids in the mutant library. B) SirT1 variants isolated after three rounds of selection from the library. C) Acyl-type preference of SirT1 variants assayed on FLuc K529ac and K529cr. D) SirT1 variant S1Ac1 deacetylates H4 K16ac but is compromised in histone decrotonylation. Activity was measured by Western blot on isolated human histones using antibodies against H4 K16ac or pan-anti-CrK.

The spectrum of isolates retrieved from different negative selections was very similar and converged on eight different sequences, whereby two of these corresponded to more than 80% of the sequenced clones. We compared the selectivity of the SirT1 mutants for their ability to deacylate FLuc K529ac and K529cr. All mutants were shifted in their substrate preference towards lysine deacetylation compared to SirT1 (Fig. 5C).

The best mutants discriminated between acetylated and crotonylated FLuc more than 20 times better than SirT1, while maintaining close to wild-type activity towards the acetylated form. This increased selectivity was maintained towards histones. SirT1Ac1 deacylated H4 K16ac with near wild-type activity but was almost 20 times less efficient in removing lysine crotonylation (Fig. 5D). The activity of the SirT1 variants decreased with increasing acyl chain length, indicating that a reduction in the volume of the substrate binding site causes the preference for acetylated lysine residues (Supplementary Fig. 9). This shows that the directed evolution approach is portable to other KDAC isoforms.

## Discussion

Based on the incorporation of acylated lysine residues in reporter enzymes. we have developed a procedure to identify a KDAC variant with the desired selectivity in a library of more than three million mutants. We screened the isolated mutants using 6-AU to modulate the stringency of the positive selection and measured the activity on acylated FLuc (37).

Sirtuins exhibit activity over a wide range of acyl chains (27), with some showing remarkable selectivity for subgroups of acylations (23, 39). We set out to create tools to modulate cellular acylation patterns to facilitate the analysis of their physiological relevance. Therefore, we have increased the selectivity of the naturally acyl-type promiscuous CobB by evolving a new conformation of the substrate binding loop. We suggest that crotonyl-lysine triggers the loop to adopt the new conformation by an induced fit mechanism (40, 41) because acetyl-lysine is unable to stabilize either loop conformation (Fig. 3C), which indicates that the apo-enzyme already exists in different conformational states.

The exchange of positions of F60 and W67 induced by the crotonyl group in CobB_ac2_ leads to the accumulation of Int. III (38), probably by preventing its hydrolysis due to increased shielding against water. This conclusion is supported by the observation that the equivalent of F60 (F33) in *Thermotoga maritima* Sir2 shields the reaction intermediates from premature hydrolysis and ADP-ribose formation (16), and that the equivalent of W67 (Y40) is predicted by MD simulations to control the access of water to the active site (42).

Our results demonstrate that the conformational rearrangement and the excessive shielding of the reaction cause inefficient hydrolysis. The residual decrotonylation activity of CobB_ac2_ may either result from a small fraction of the enzyme following the regular catalytic pathway or from eventual hydrolysis of Int. III. We favour the former explanation because of the longevity of the crotonyl Int. III in our structure (100% occupancy remaining after 36 h soaking) and the conformational flexibility of the CobB_ac2_-H4 K16cr complex (Supplementary Fig. 8).

The residue I131 is part of the conserved NID motif (43) of Sirtuins. Interestingly, its mutation to cysteine is present in human SirT7. The A76G mutation, to our knowledge, does not exist in other proteins. However, at the equivalent position in SirT3 the mutation F180L was found to increase selectivity for acetyl by 20-fold over crotonyl-lysine, possibly by removing the π-π stacking interaction with the crotonyl group (24). Unfortunately, these findings are not readily portable to other sirtuins because the phenylalanine residue is not fully conserved.

The SirT1 mutants appear to have acquired selectivity by a different mechanism than CobB_ac2_, since they show a marked decrease in activity with increasing acyl chain length. All mutants show a redistribution of hydrophobic residues, indicating a change of selectivity due to alternative hydrophobic packing of the substrate binding pocket. Similar results were obtained by engineering SH3 domains (44), ubiquitin (45) and human carbonic anhydrase II (46).

Acyl-type specific variants of KDACs will be useful to deconvolute the individual contributions of different types of lysine acylations to cell physiology (11). They can be employed to erase a particular acylation by overexpression in the wild-type background or to partially complement knockout cell lines. Targeted to specific genomic regions, they can be used to locally deplete particular types of acylation and to investigate the impact of metabolites on gene expression (47).

Rational design of acyl-type selective mutants is challenging and often results in partial success. CobB mutants with up to 43-fold increased preference for AcK over N(ε)-succinyl-lysine have been created in this way (48). A crotonyl-lysine selective HDAC1 was used to investigate the role of this modification in transcriptional regulation and self-renewal of mouse embryonic stem cells (20). In these cases, selectivity was successfully engineered by removing acyl-specific interactions. Obviously, however, there is a more complex interaction between the active site residues in sirtuins, making a rational design of selective variants very difficult. Thus, our selection system is a powerful tool to find such KDAC mutants which will be tremendously useful for future mechanistic studies.

With the selection system described here, KDAC variants are easily identifiable for any type of lysine modification accessible by genetic code expansion. Furthermore, the selection system could also be used to facilitate the creation of selective mutant/inhibitor pairs by a bump-and-hole strategy (49) and may allow the creation of enzymes with novel bioorthogonal reactivity.

## Materials and Methods

### Demodification reactions of myoglobin K4mod

Modified myoglobin K4_mod_ was expressed and purified according to published protocols (32). The demodification reaction was setup in a 50 μL volume containing: 40 mM Tris pH 8, 50 mM NaCl, 1 mM DTT, 0.1 mM ZnCl_2_, 3 mM NAD^+^, 30 μM myoglobin K4_mod_ and purified CobB (4 μM wildtype, 16 μM CobB_ac2_, 48 μM CobB_ac3_ and 36 μM CobB_ac6_). The reactions were incubated at 30°C for 16 h and stopped by chilling at 4°C. 12 μL of each reaction was injected on a HPLC (Agilent 1100, Agilent Technologies) and desalted by C4 column. The proteins were eluted with a gradient of 20-80% acetonitrile in water (0.1% TFA) and analyzed by ESI-MS (LCQ Advantage MAX, Thermo Finnigan). Protein mass was calculated by MagTran and myoglobin peaks were normalized on maximal peak intensity (50).

### Purification of Firefly Luciferase K529_mod_

*E. coli* BL21 DE3 RIL were transformed with plasmids pCDF-PylT-FLuc(opt)His6-K529TAG and pBK-AcKRS3opt (for FLuc K529ac) or pBK-PylS (51) (other modifications). Cells were grown in LB medium in the presence of antibiotics (50 μg/ml spectinomycin and 50 μg/ml kanamycin) to an OD_600_ of 0.3 at 37°C. Then, cells were shifted to 30°C, after 1 h protein expression was induced by the addition of 1 mM IPTG, modified-lysine (acetyl-lysine 10 mM, others 1 mM) and 50 mM NAM. After 16 h at 30°C cells were harvested by centrifugation, washed with PBS and suspended in Ni-wash buffer (20 mM Tris/HCl pH 8, 10 mM imidazole, 200 mM NaCl, 10 mM DTT, 2 mM PMSF, 0.5x Roche Protease Inhibitor cocktail, containing 20 mM NAM and lysozyme. Lysis was preformed using a pneumatic cell disintegrator the cell debris removed by centrifuged (20 min, 50,000 g, 4°C). The supernatant was supplemented with 500 μl Ni-NTA-beads. After 2 h incubation with agitation at 4°C beads were washed with 30 ml Ni-NTA wash buffer and bound proteins eluted in Ni-NTA wash buffer supplemented with 200 mM imidazole. The eluate buffer was exchanged (20 mM Tris pH 8, 50 mM NaCl, 10 mM DTT) and used in appropriate dilution (s. below) as substrate for the KDAC assays.

### Luciferase-based KDAC assay

Typical endpoint deacetylation reactions contain: 5 μL diluted Firefly Luciferase K529mod (Final dilution: 1:1500 Ac, 1:500 Bu, 1:125 Cr and Pr), 2 mM NAD^+^, 2 μM KDAC in 50 μl KDAC buffer (25 mM Tris/HCl pH 8.0, 137 mM NaCl, 2.7 mM KCl, 1 mM MgCl_2_, 1 mM DTT, 1 mg/ml BSA). The reactions are incubated for 2 h at 30°C. Luciferase activity is then assayed by addition of an equal volume of a mixture containing 40 mM Tricine pH 7.8, 200 μM EDTA, 7.4 mM MgSO_4_, 2 mM NaHCO_3_, 34 mM DTT, 0.5 mM ATP and 0.5 mM luciferin) (52). Luminescence is quantified using a FluoStar Omega Microplate Reader (BMG Labtech).

### Acidic extract of histone proteins from HEK293 GnTI^−^ cells

HEK293 GnTI^−^ cells were grown in Freestyle medium supplemented with 2% FBS in an orbital shaker at 37°C with 8% CO_2_ to a density of 2-3 × 106 cells/ml. The cells were incubated for 8 h at 37°C with 8% CO_2_ before addition of 10 mM sodium butyrate to the flask. Following the addition, the flask were shifted to 30°C with 8% CO2 while shaking at 130 rpm. Cells were harvested 40 h after addition of butyrate. Cells were suspended in lysis buffer (25 mM Tris pH 8, 300 mM NaCl, 10% glycerol, 1 mM TCEP, 1 mM EDTA, 5 μg/ml AEBSF, 5 μg/ml aprotinin, 5 μg/ml leupeptin) and lysed by passing through a microfluidizer. Cell debris were collected by centrifugation at 15.000 g for 5 min at 4°C. The nuclear histones were extracted from the cell debris following the acidic extraction protocol of Sechter et al. (53). After TCA precipitation the histone protein was suspended in water at a concentration of 1 mg/mL and used as substrate stock.

### Sirtuin reaction on histone extract and Western blot analysis

Each reaction contained 250 ng histone extract, Sirtuin (64 nM to 4 μM) and 2 mM NAD^+^ in KDAC buffer (25 mM Tris/HCl pH 8.0, 137 mM NaCl, 2.7 mM KCl, 1 mM MgCl_2_, 1 mM DTT, 1 mg/ml BSA). The reaction was incubated for 2 h at 30°C and stopped by adding 10 μL 3x SDS-Sample Buffer. Samples were mixed and heated to 90°C for 10 min. 5 μL of the reaction mixture were resolved by 15% SDS-PAGE and histone modifications detected by Western blot. The H4 K16ac modification was detected using a monoclonal antibody (Abcam, ab109463) and lysine crotonylation was detected using a pan-specific anti-CrK antibody (PTM Biolabs, PTM105), both were used according to manufacturer’s specifications.

### KDAC selection system

To test the selection strategy for CobB (Fig. 1B), *E. coli* DB6656 (Δ*pyrF*) were transformed with pPylT-URA3, pPylT-URA3-K93TAG-PylS or pPylT-URA3-K93TAG-AcKRS3 and grown overnight in LB-medium (15 μg/mL tetracycline). The cells were replicated in tenfold dilutions with M9 medium (100 to 10-7) onto M9 selection plates (M9 minimal medium, 0.2% arabinose, 1% glycerol, 0.1 mM tryptophan, 50 μg/mL kanamycin, 15 μg/mL tetracycline, ± 0.1 mM uracil, ± 10 mM acetyl-lysine and 1 mM boc-lysine, ± 0.1% 5-fluoroorotic acid). Plates were imaged after incubation for 48 h at 37°C.

To adapt the selection system to HDAC8, SirT1, SirT2 and SirT3, *E. coli* DH10B Δ*pyrF* Δ*cobB* was transformed in two sequential steps first with either pPylT-URA3, pPylT-URA3-K93TAG-PylS or pPylT-URA3-K93TAG-AcKRS3 and next with either pBK-His_6_-HDAC8, pBK-His_6_-SirT1cat, pBK-His_6_-TEV-SirT2cat or pBK-His_6_-TEV-SirT3cat. Transformants were grown overnight in 4 mL LB (50 μg/mL kanamycin, 15 μg/mL tetracycline) and replicated in a tenfold dilution series onto M9 selection plates (M9 minimal medium, 0.2% arabinose, 1% glycerol, 0.4% glucose, 0.1 mM tryptophan, 80 mg/L valine, 80 mg/L isoleucine, 80 mg/L leucine, 50 μg/mL kanamycin, 15 μg/mL tetracycline, ± 0.1 mM uracil, ± 10 mM acetyl-lysine and 1 mM boc-lysine, ± 0.1% 5-fluoroorotic acid). Plates were imaged after incubation for 48 h at 37°C.

### CobB active site library creation

The active site mutant library was designed based on the CobB crystal structure 1S5P (54). The codons for A76, Y92, R95, I131, V187 were replaced by NNK codons in three rounds of inverse PCR using the Expand™ High Fidelity PCR System (Roche). One PCR reaction contained 1x Buffer 2, 0.2 mM dNTPs, 1 μM forward and reverse primer, 100 ng pBK-CobB plasmid DNA, 3.5 U Expand Polymerase in a 50 μL volume and run with the following program:

- 95° 1 min
- 95° 20 s 10x
- 65° 30 10x −1°C/cycle
- 68° 5 min 10x repeat cycle
- 95° 20 s 25x
- 55° 30 25x
- 68° 5 min 25x repeat cycle
- 68° 5 min
- 4° hold

The PCR reaction was purified using the QIAquick PCR Purification Kit (Qiagen) after digestion of the template plasmid with 20 U Dpn I (NEB) for 1 h at 37°C. The purified reaction was digested with 80 U HF-Bsa I (NEB) for 2 h and purified as before. The digested PCR product was circularized by T4 DNA ligase in a total volume of 500 μL 1x T4 DNA Ligase Reaction Buffer containing 100 μL digested PCR product and 10 kU T4 DNA Ligase (NEB). The reaction was incubated overnight at 16°C and purified by ethanol precipitation. The precipitated DNA was dissolved in 10 μL water and used to electroporate MAX Efficiency™ DH10B™ Competent Cells (ThermoFisher) according to the manufacturer's specifications. Transformation efficiency was determined from the colony count of a dilution series (10^−4^ to 10^−7^) on LB agar plates (50 μg/mL kanamycin) for every round of mutagenesis. Coverage was calculated using the GLUE-IT web tool and was estimated to cover for 99.6% of all possible amino acid combinations in the final round with 108 transformants (55). The plasmid DNA was isolated from the mutant pool by Plasmid Maxi Kit (QIAGEN) according to the manufacturer's specifications.

### Library creation of SirT1

The SirT1 mutant library was created in the same manner as the CobB library. Positions A313, I316, I347, F366L and I411 were mutated to NNK in 3 rounds of mutagenesis. The final transformation step had a transformation efficiency of 1.8 × 10^7^ covering 85% of all possible variants (55). The plasmid DNA was isolated from the mutant pool by Plasmid Maxi Kit (QIAGEN) according to the manufacturer's specifications.

### Library selection

Freshly prepared electrocompetent *E. coli* DH10B Δ*pyrF* Δ*cobB* containing pPylT-URA3-K93TAG-PylS or pPylT-URA3-K93TAG-AcKRS3 were electroporated with the CobB or SirT1 mutant library with an efficiency to cover >95% of the diversity and incubated at 37°C overnight. Before plating the cells, the pool was diluted 1:50 in 50 mL LB (50 μg/mL kanamycin, 15 μg/mL tetracycline) and incubated for 3 h at 37°C and 200 rpm. After 3 h the cells were pre-incubated with the amino acid (10 mM acetyl-lysine or 1 mM for other modifications) and 0.2% arabinose. After 3 h, 1 mL cells were harvested by centrifugation (8000 rpm, 2 min) and washed twice with 1 mL PBS to remove the amino acid and uracil. The washed cells were resuspended to an OD_600_ of 1 in PBS. 100 μL cell suspension was plated on the selection plate (positive selection: M9 selection plate +amino acid, –uracil, –5-FOA or negative selection: M9 selection plate +amino acid, +uracil, +5-FOA) and a control plate (M9 selection plate, –amino acid, –uracil, –5-FOA). The plates were incubated for at least 48 h at 37°C. Colonies were scraped from the plate and plasmid DNA isolated with GeneJET Plasmid Miniprep Kit (Thermo Scientific). The isolated plasmid DNA was used to transform *E. coli* cells for subsequent selections. To obtain CobB mutants with preference for deacetylation over decrotonylation, three rounds of selection on the active site library were performed, starting with a positive selection for deacetylation followed by negative selection against decrotonylation and a final positive selection for deacetylation. 72 single colonies of the last selection were picked from the selection plate, grown in LB (1 mL, 50 μg/mL kanamycin), plasmid DNA isolated, retransformed in DH10B by heat shock, plasmid DNA isolated and sequenced.

### Activity plate assay using 6-Azauridine (6-AU)

Individual plasmid isolates were retransformed in *E. coli* DH10B Δ*pyrF* Δ*cobB* containing pPylT-URA3-K93TAG-AcKRS3 and grown in 1 mL LB (50 μg/mL kanamycin, 15 μg/mL tetracycline) in a 96 well block. Cells were replicated using a pin head replicator onto LB agar plates (50 μg/mL kanamycin, 15 μg/mL tetracycline), M9 selection plates and selection plates containing increasing amounts of 6-AU (positive selection: M9 selection plate +Acetyl-Lysine, –uracil, –5-FOA, + 0.01-2 mM 6-AU). Cells were imaged after 48 h.

## Acknowledgment

We thank Petra Geue for technical support and Sascha Gentz for peptide synthesis. Datasets were collected at PXII X10SA beamline at the Suisse Light Source, Villigen, Switzerland. We thank local contacts and beamline scientists for their support. This project was supported financially by the Heisenberg Program of the German Research Foundation (DFG) [NE1589/5-1 to H.N.] and the Max-Planck-Institute of Molecular Physiology.

## References

1. Allfrey VG, Faulkner R, & Mirsky AE (1964) Acetylation and Methylation of Histones and Their Possible Role in the Regulation of Rna Synthesis. Proc Natl Acad Sci U S A 51:786–794.

2. Verdin E & Ott M (2015) 50 years of protein acetylation: from gene regulation to epigenetics, metabolism and beyond. Nat Rev Mol Cell Biol 16(4):258–264.

3. Feige JN & Auwerx J (2008) Transcriptional targets of sirtuins in the coordination of mammalian physiology. Current Opinion in Cell Biology 20(3):303–309.

4. Wilkins BJ, et al. (2014) A Cascade of Histone Modifications Induces Chromatin Condensation in Mitosis. Science (New York, N.Y.) 343(6166):77.

5. Tas RP, et al. (2017) Differentiation between Oppositely Oriented Microtubules Controls Polarized Neuronal Transport. Neuron 96(6):1264–1271.e1265.

6. Houtkooper RH, Pirinen E, & Auwerx J (2012) Sirtuins as regulators of metabolism and healthspan. Nature Reviews Molecular Cell Biology 13:225.

7. Jing E, et al. (2011) Sirtuin-3 (Sirt3) regulates skeletal muscle metabolism and insulin signaling via altered mitochondrial oxidation and reactive oxygen species production. Proceedings of the National Academy of Sciences 108(35):14608.

8. Haberland M, Johnson A, Mokalled MH, Montgomery RL, & Olson EN (2009) Genetic dissection of histone deacetylase requirement in tumor cells. Proceedings of the National Academy of Sciences 106(19):7751.

9. Stilling RM & Fischer A (2011) The role of histone acetylation in age-associated memory impairment and Alzheimer’s disease. Neurobiology of learning and memory 96(1):19–26.

10. Rogina B & Helfand SL (2004) Sir2 mediates longevity in the fly through a pathway related to calorie restriction. Proceedings of the National Academy of Sciences of the United States of America 101(45):15998.

11. Choudhary C, Weinert BT, Nishida Y, Verdin E, & Mann M (2014) The growing landscape of lysine acetylation links metabolism and cell signalling. Nat Rev Mol Cell Biol 15(8):536–550.

12. Lin H, Su X, & He B (2012) Protein lysine acylation and cysteine succination by intermediates of energy metabolism. ACS Chem Biol 7(6):947–960.

13. Yang XJ & Seto E (2008) The Rpd3/Hda1 family of lysine deacetylases: from bacteria and yeast to mice and men. Nat Rev Mol Cell Biol 9(3):206–218.

14. Feldman JL, Dittenhafer-Reed KE, & Denu JM (2012) Sirtuin catalysis and regulation. J Biol Chem 287(51):42419–42427.

15. Sanders BD, Jackson B, & Marmorstein R (2010) Structural basis for sirtuin function: what we know and what we don’t. Biochim Biophys Acta 1804(8):1604–1616.

16. Hawse WF, et al. (2008) Structural Insights into Intermediate Steps in the Sir2 Deacetylation Reaction. Structure 16(9):1368–1377.

17. Landry J, et al. (2000) The silencing protein SIR2 and its homologs are NAD-dependent protein deacetylases. Proceedings of the National Academy of Sciences 97(11):5807.

18. Tanner KG, Landry J, Sternglanz R, & Denu JM (2000) Silent information regulator 2 family of NAD- dependent histone/protein deacetylases generates a unique product, 1-*O*-acetyl-ADP-ribose. Proceedings of the National Academy of Sciences 97(26):14178.

19. Zhao S, Zhang X, & Li H (2018) Beyond histone acetylation—writing and erasing histone acylations. Current Opinion in Structural Biology 53:169–177.

20. Wei W, et al. (2017) Class I histone deacetylases are major histone decrotonylases: evidence for critical and broad function of histone crotonylation in transcription. Cell research 27(7):898–915.

21. Feldman JL, Baeza J, & Denu JM (2013) Activation of the protein deacetylase SIRT6 by long-chain fatty acids and widespread deacylation by mammalian sirtuins. The Journal of biological chemistry 288(43):31350–31356.

22. Tan M, et al. (2014) Lysine glutarylation is a protein posttranslational modification regulated by SIRT5. Cell metabolism 19(4):605–617.

23. Du J, et al. (2011) Sirt5 is a NAD-dependent protein lysine demalonylase and desuccinylase. Science (New York, N.Y.) 334(6057):806–809.

24. Bao X, et al. (2014) Identification of ‘erasers’ for lysine crotonylated histone marks using a chemical proteomics approach. eLife 3:e02999.

25. Cao J, et al. (2019) HDAC11 regulates type I interferon signaling through defatty-acylation of SHMT2. Proceedings of the National Academy of Sciences 116(12):5487.

26. Sauve AA (2010) Sirtuin chemical mechanisms. Biochimica et biophysica acta 1804(8):1591–1603.

27. Bheda P, Jing H, Wolberger C, & Lin H (2016) The Substrate Specificity of Sirtuins. Annual Review of Biochemistry 85(1):405–429.

28. Zhao K, Harshaw R, Chai X, & Marmorstein R (2004) Structural basis for nicotinamide cleavage and ADP-ribose transfer by NAD^+^-dependent Sir2 histone/protein deacetylases. Proceedings of the National Academy of Sciences of the United States of America 101(23):8563–8568.

29. Neumann-Staubitz P & Neumann H (2016) The use of unnatural amino acids to study and engineer protein function. Curr Opin Struct Biol 38:119–128.

30. Miller BG, Snider MJ, Wolfenden R, & Short SA (2001) Dissecting a charged network at the active site of orotidine-5’-phosphate decarboxylase. J Biol Chem 276(18):15174–15176.

31. Neumann H & Chin J (2009) Methods and compositions. (Google Patents).

32. Neumann H, Peak-Chew SY, & Chin JW (2008) Genetically encoding N(epsilon)-acetyllysine in recombinant proteins. Nat Chem Biol 4(4):232–234.

33. Meng X, Smith RM, Giesecke AV, Joung JK, & Wolfe SA (2006) Counter-selectable marker for bacterial-based interaction trap systems. Biotechniques 40(2):179–184.

34. Gattner MJ, Vrabel M, & Carell T (2013) Synthesis of epsilon-N-propionyl-, epsilon-N-butyryl-, and epsilon-N-crotonyl-lysine containing histone H3 using the pyrrolysine system. Chem Commun (Camb) 49(4):379–381.

35. Kim CH, Kang M, Kim HJ, Chatterjee A, & Schultz PG (2012) Site-specific incorporation of epsilon-N-crotonyllysine into histones. Angew Chem Int Ed Engl 51(29):7246–7249.

36. Wilkins BJ, et al. (2015) Genetically encoding lysine modifications on histone H4. ACS Chem Biol 10(4):939–944.

37. Spinck M, Ecke M, Sievers S, & Neumann H (2018) Highly Sensitive Lysine Deacetylase Assay Based on Acetylated Firefly Luciferase. Biochemistry 57(26):3552–3555.

38. Wang Y, et al. (2017) Deacylation Mechanism by SIRT2 Revealed in the 1′-SH-2′-O-Myristoyl Intermediate Structure. Cell Chemical Biology 24(3):339–345.

39. Teng Y-B, et al. (2015) Efficient Demyristoylase Activity of SIRT2 Revealed by Kinetic and Structural Studies. Scientific reports 5:8529.

40. Bosshard HR (2001) Molecular Recognition by Induced Fit: How Fit is the Concept? Physiology 16(4):171–173.

41. Koshland DE (1958) Application of a Theory of Enzyme Specificity to Protein Synthesis. Proceedings of the National Academy of Sciences 44(2):98–104.

42. Shi Y, Zhou Y, Wang S, & Zhang Y (2013) Sirtuin Deacetylation Mechanism and Catalytic Role of the Dynamic Cofactor Binding Loop. The journal of physical chemistry letters 4(3):491–495.

43. Brachmann CB, et al. (1995) The SIR2 gene family, conserved from bacteria to humans, functions in silencing, cell cycle progression, and chromosome stability. Genes & development 9(23):2888–2902.

44. Ben-David M, et al. (2019) Allosteric Modulation of Binding Specificity by Alternative Packing of Protein Cores. J Mol Biol 431(2):336–350.

45. Haririnia A, et al. (2008) Mutations in the Hydrophobic Core of Ubiquitin Differentially Affect Its Recognition by Receptor Proteins. J Mol Biol 375(4):979–996.

46. Hunt JA, Ahmed M, & Fierke CA (1999) Metal Binding Specificity in Carbonic Anhydrase Is Influenced by Conserved Hydrophobic Core Residues. Biochemistry 38(28):9054–9062.

47. Heitmuller S, Neumann-Staubitz P, Herrfurth C, Feussner I, & Neumann H (2018) Cellular substrate limitations of lysine acetylation turnover by sirtuins investigated with engineered futile cycle enzymes. Metab Eng 47:453–462.

48. Colak G, et al. (2013) Identification of lysine succinylation substrates and the succinylation regulatory enzyme CobB in Escherichia coli. Mol Cell Proteomics 12(12):3509–3520.

49. Shah K, Liu Y, Deirmengian C, & Shokat KM (1997) Engineering unnatural nucleotide specificity for Rous sarcoma virus tyrosine kinase to uniquely label its direct substrates. Proc Natl Acad Sci U S A 94(8):3565–3570.

50. Zhang Z & Marshall AG (1998) A universal algorithm for fast and automated charge state deconvolution of electrospray mass-to-charge ratio spectra. Journal of the American Society for Mass Spectrometry 9(3):225–233.

51. Neumann H, Peak-Chew SY, & Chin JW (2008) Genetically encoding Nε-acetyllysine in recombinant proteins. Nature chemical biology 4:232.

52. Siebring-van Olst E, et al. (2013) Affordable luciferase reporter assay for cell-based high-throughput screening. J Biomol Screen 18(4):453–461.

53. Shechter D, Dormann HL, Allis CD, & Hake SB (2007) Extraction, purification and analysis of histones. Nature Protocols 2:1445.

54. Zhao K, Chai X, & Marmorstein R (2004) Structure and substrate binding properties of cobB, a Sir2 homolog protein deacetylase from Escherichia coli. J Mol Biol 337(3):731–741.

55. Firth AE & Patrick WM (2008) GLUE-IT and PEDEL-AA: new programmes for analyzing protein diversity in randomized libraries. Nucleic Acids Research 36(Web Server issue):W281–W285.

