## Supplementary Methods and Figures for "Evolved, Selective Erasers of Distinct Lysine Acylations"

### Directed evolution of lysine deacetylases

### Supplementary methods

*Plasmids*

pPylT-URA3

pMyo4TAG-PylT-based plasmid(1) contains p15A origin, tetracycline resistance, PylT of *Methanosarcina barkeri* and URA3 under arabinose promotor inserted by Nco I/Xho I restriction sites.

pPylT-URA3-K93TAG-PylS, pPylT-URA3-K93TAG-AcKRS3

Based on pPylT-URA3 plasmid. TAG codon was introduced at the position of K93 by QuickChange mutagenesis. Either the *Methanosarcina barkeri* PylS or AcKRS3 (2) were introduced downstream of URA3 via XhoI/KpnI restriction sites.

pCDF-PylT-FLuc(opt)His6-K529TAG

The gene for Firefly Luciferase codon-optimized for expression in *E. coli* and containing an amber codon replacing the codon for Lys-529 as well as a C-terminal $His_6$-Tag was custom synthesized by Genscript and cloned into Nco I/Xho I of pCDF-PylT (3).

pBK-AcKRS3opt

(expressing acetyl-lysyl-tRNA synthetase with mutations improving tRNA binding) was generated from pBK-AcKRS3 by three rounds of QuickChange mutagenesis introducing mutations V31I, T56P, H62Y and A100E (4).

pBK-$His_6$-CobB

$His_6$-CobB was amplified from pBAD-CobB-H3-H4-Hat1 (5) with Bgl II/Stu I sites and cloned into BamH I/Stu I sites of pBK-PylS (1).

pBK-His$_6$-HDAC8

His$_6$-HDAC8 gene was custom synthesized by Invitrogen. It was amplified from pMA-T backbone with Nco I/Xba I sites and cloned sticky/blunt ended into Nco I/Stu I sites downstream of the Ara-BAD promotor of pBK-CobB, replacing the CobB gene.

pBK-His$_6$-SirT1cat

His$_6$-SirT1cat (encoding aa 225-664 of SirT1) was amplified from pRSF-Duet1-His6-SirT1 (6) with Nco I/Xba I sites and cloned into Nco I and Xba I sites of pBK-His6-hsHDAC8.

pBK-His$_6$-TEV-SirT2cat and pBK-His$_6$-TEV-SirT3cat

The catalytic domain of SirT2 (aa 56-356) and SirT3 (aa 118-399) were amplified from pGEX-T5S-TEV-SirT2/3 (6) with Nco I/Xba I sites, His$_6$-Tag and TEV protease cleavage site and cloned into pBK-His6-hsHDAC8 using the Nco I and Xba I sites.

pCDF-His$_6$-Sirt1NCatC

For expression, the mutations were transferred from pBK-His$_6$-Sirt1cat to pCDF-His$_6$-Sirt1NCatC containing a N and C-terminal extended SirT1 (158-665). The mutations were transferred using restriction cloning in BglII and HindIII sites.

pCRISPR-CobB-PAM, pCRISPR_spec-PyrF-PAM

New spacer sequences were introduced in pCRISPR (#Addgene 42875) using the manufacture protocol. As CobB spacer, the following oligos were used 5'-aaa cTG GAA AAA CCA AGA GTA CTC GTA CTG ACA Gg-3' and 5'-aaa acC TGT CAG TAC GAG TAC TCT TGG TTT TTC CA-3'.

For PyrF the oligos 5’-AAA CAA GGT CGG CAA AGA GAT GTT TAC ATT GTT TG-3’ and 5’-AAA ACA AAC AAT GTA AAC ATC TCT TTG CCG ACC TT-3’ were used. Also, the antibiotic resistance marker of the plasmid was changed to spectinomycin by restriction cloning using Xho I/Spe I sites.

*Strains*

DB6656 was obtained from CGSC (CGSC#: 6868), DH10B was purchased from ThermoFisher. DH10B *ΔpyrF ΔcobB* was created using CRISPR/Cas9: The genes encoding PyrF and CobB were inactivated in *E. coli* DH10B by inserting several stop codons shortly after the start codon of the respective gene in two sequential rounds of genome engineering using CRISPR/Cas9 and editing oligos 5’-CAC GCT GTT GAA GTT CGC GCA CAA ACT GTT AAG CTT ACA ATG TAA ACA TCT CTT TGC CGA GGT TC-3’ (pyrF) and 5’-AGG TAC GAA TAC CTG ATT CCG CAG AAA TTt aag ctt aTG TCA GTA CGA GTA CTC TTG GTT TTT CCA-3’ (cobB) following established protocols (7). Successful mutagenesis was confirmed by colony PCR and restriction digest analysis of a simultaneously introduced Hind III site. Plasmids were cured with 60 μg/mL promazine (8).

*Expression of CobB*

*E. coli* DH10B Δ*pyrF* Δ*cobB* pPylT-URA3 was transformed with the respective pBK plasmids for CobB wild-type or mutant. An overnight culture (ONC) in 10 mL LB-medium (50 μg/mL kanamycin, 15 μg/mL tetracycline) was prepared. The ONC was used to inoculate 1 L LB medium (50 μg/mL kanamycin, 15 μg/mL tetracycline) and cells were grown to an $OD_{600}$ of 0.3. The temperature was reduced to 30°C 1 h before expression was induced by addition of arabinose to a final concentration of 0.2%. Cells were harvested after 16 h by centrifugation (20 min, 6000 rpm, 4°C). The cell pellets were washed with PBS and stored at –20°C.

*Expression of SirT1*

BL21(DE3) cells were transformed with wild-type or mutant pCDF-$His_6$-SirT1-NcatC plasmid. An overnight culture was prepared in 10 mL LB (50 µg/mL spectinomycin) and incubated at 37°C while agitating at 200 rpm. The expression culture was inoculated with the preculture and grown at 37°C and 200 rpm to an $OD_{600}$ of 0.8. The expression was induced by addition of 0.5 mM IPTG and the culture shifted to 18°C. The cells were harvested by centrifugation the next day, washed with PBS and stored at –20°C.

*Purification of KDACs*

a) CobB variants: Cell pellets were thawed on ice and resuspended in 25 mL Ni-NTA wash buffer (20 mM HEPES pH 7.5, 200 mM NaCl, 20 mM imidazole, 1 mM DTT) supplemented with lysozyme (~0.5 mg/mL), DNase (1 mg) and protease inhibitor (1 mM PMSF). Lysis was preformed using a pneumatic cell disintegrator. The cell debris was removed by centrifugation (20 min, 20.000 rpm, 4°C) and HisPur™ $Ni^{2+}$-NTA resin (2 mL in 50 mL solution) was added to the supernatant. After 1 h at 4°C the suspension was loaded on a plastic column (BioRad, Munich) with a frit and washed with 40 mL Ni-NTA wash buffer. Protein was eluted in 4 mL Ni-NTA wash buffer supplemented with 200 mM imidazole. The elutate was concentrated and the buffer exchanged to gelfiltration buffer

(20 mM HEPES pH 7.5, 100 mM NaCl, 10 mM DTT) before loading on a HILoad™ 26/70 Superdex™ 200 size-exclusion chromatography column (GE healthcare, UK) preequilibrated with gel filtration buffer. Absorption at 280 nm was monitored and 5 mL fractions collected. Fractions containing protein were analyzed on a SDS-PAGE, pooled and concentrated in a microfiltrator (Amicon Ultra-15 Centrifugal Unit, 10 kDa, Merck Millipore). The protein was aliquoted (50 μL), flash frozen in liquid nitrogen and stored at –80°C.

b) SirT1 variants: The cell pellet was resuspended in wash buffer (20 mM Tris pH 8, 200 mM NaCl, 20 mM Imidazole) supplemented with 0.2 mM PMSF, lysozyme (0.5 mg/mL) and DNAse (ca 1 mg). The cells were lysed by passing through a microfuidizer and cleared from debris by centrifugation (20 min, 20.000 g). The supernatant was mixed with 500 μL HisPur™ $Ni^{2+}$-NTA resin and agitated for 1.5 h at 4°C. The suspension was filtered through a plastic column (BioRad, Munich) with a frit and the beads washed with 40 volumes wash buffer. The protein was eluted from the beads with 2 mL wash buffer supplemented with 200 mM imidazole. The protein was concentrated and the buffer exchanged with gel filtration buffer (20 mM Tris pH 8, 50 mM NaCl, 10 mM DTT). The protein was applied to a Superdex 75 gel filtration column; 1 mL fractions were collected and analyzed on an SDS-PAGE. Fractions containing SirT1 were concentrated to about 10 mg/mL, flash frozen in liquid nitrogen and stored at –80°C.

*Acidic extract of histone proteins from HEK293 GnTI⁻ cells*

HEK293 $GnTI^-$ cells were grown in Freestyle medium supplemented with 2% FBS in an orbital shaker at 37°C with 8% $CO_2$ to a density of 2-3 × $10^6$ cells/ml. The cells were incubated for 8 h at 37°C with 8% $CO_2$ before addition of 10 mM sodium butyrate to the flask. Following the addition, the flask were shifted to 30°C with 8% $CO_2$ while shaking at 130 rpm. Cells were harvested 40 h after addition of butyrate. Cells were suspended in lysis buffer (25 mM Tris pH 8, 300 mM NaCl, 10% glycerol, 1 mM TCEP, 1 mM EDTA, 5 µg/ml AEBSF, 5 µg/ml aprotinin, 5 µg/ml leupeptin) and lysed by passing through a microfluidizer. Cell debris were collected by centrifugation at 15.000 g for 5 min at 4°C. The nuclear histones were extracted from the cell debris following the acidic extraction protocol of Sechter et al. (9). After TCA precipitation the histone protein was suspended in water at a concentration of 1 mg/mL and used as substrate stock.

*Selectivity assay using Ni-Eluate of KDAC variants.*

Individual plasmid isolates were expressed and purified as previously described. The buffer of the Ni-NTA eluate was exchanged to gel filtration buffer, purity and concentration were approximated by SDS-PAGE. Luciferase-based KDAC reactions were prepared with an approximated equal concentration (2 µM final, verified on SDS-PAGE) of each mutant for different modified Fluc substrates (Ac, Bu, Cr, Pr) and relative mutant activity for each substrate was normalized to the wt activity.

*Crystallization of CobB mutants*

The crystallization of CobB, $CobB_{ac2}$ and $CobB_{ac3}$ were set up as reported for CobB (10). Crystals were obtained for CobB after 1 day from hanging drop crystallization with 100 mM Bis-Tris, 30 mM HCl, 22% PEG 3350 as precipitant and 10 mg/mL $CobB_{ac2}$ preincubated with 1 mM H4 $K16_X$ (X: Ac, Bu, Cr) peptide. Crystals were flash frozen in liquid nitrogen with 20% PEG400 as cryoprotectant containing 1 mM H4 $K16_X$ (X: Ac, Bu, Cr).

Crystals were obtained for $CobB_{ac2}$ after 1 day from hanging drop crystallization with 100 mM Bis-Tris, 30 mM HCl, 23% PEG 3350 (Bu: 24%) as precipitant and 10.5 mg/mL $CobB_{ac2}$ preincubated with 1 mM H4 $K16_X$ (X: Ac, Bu, Cr) peptide. Crystals were flash frozen in liquid nitrogen with 20% PEG400 as cryoprotectant containing 1 mM H4 $K16_X$ (X: Ac, Bu, Cr).

Crystals from $CobB_{ac3}$ were obtained after 1 week from hanging drop crystallization with 100 mM Bis-Tris, 10 mM HCl, 22% PEG 3350 as precipitant and 7.2 mg/mL $CobB_{ac3}$ preincubated with 1 mM H4 $K16_{Ac}$ peptide. Crystals were flash frozen in liquid nitrogen with 20% glycerol as cryoprotectant containing 1 mM H4 $K16_{Ac}$.

The $CobB_{ac2}$-H4$K16_{Cr}$ -$NAD^+$ complex was obtained after 16 h hanging drop crystallization with 100 mM Bis-Tris, 20 mM HCl, 22% PEG 3350 as precipitant and 11.1 mg/mL $CobB_{ac2}$ preincubated on ice with 1 mM $NAD^+$ and 1 mM H4 $K16_{Cr}$ peptide. Crystals were flash frozen in liquid nitrogen with 20% PEG400 as cryoprotectant containing 1 mM $NAD^+$ and 1 mM H4 $K16_{Cr}$.

For soaking the initial crystallization of $CobB_{Ac2}$-H4$K16_{Cr}$ was replicated and crystals obtained. Crystals were transferred using a nylon loop in a 1 uL hanging drop supplemented with 1 mM H4$K16_{Cr}$ peptide and 1 mM $NAD^+$ and incubated for 2 h and 36 h. Crystals were flash frozen in liquid nitrogen and 20% PEG400 as cryoprotectant containing 1 mM $NAD^+$ and 1 mM H4 $K16_{Cr.}$

Data collection was performed at beamline PXII X10SA at the Suisse Light Source, Villigen, Switzerland. The data was processed with XDS and scaled using XSCALE (11).

Molecular replacement was performed with Phaser (12) and the data was refined using Refmac5 (13) initially and Phenix.refine (14) in the final steps. The ligand geometry file was created using JLigand (15) for coordinates and PRODRG (16) for calculation of restraints. Model building was done with Coot (17).

### Supplementary Figures

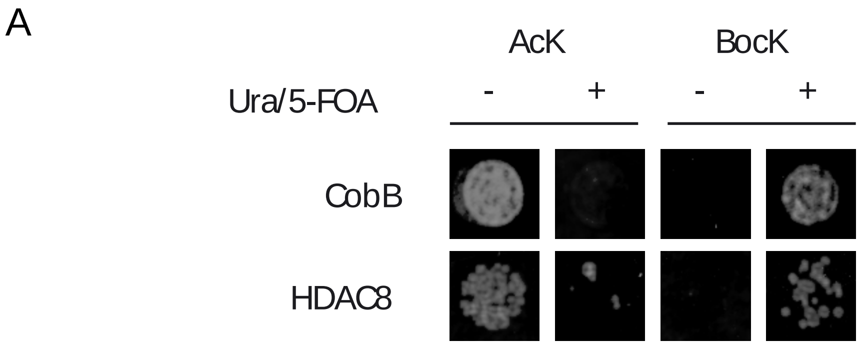

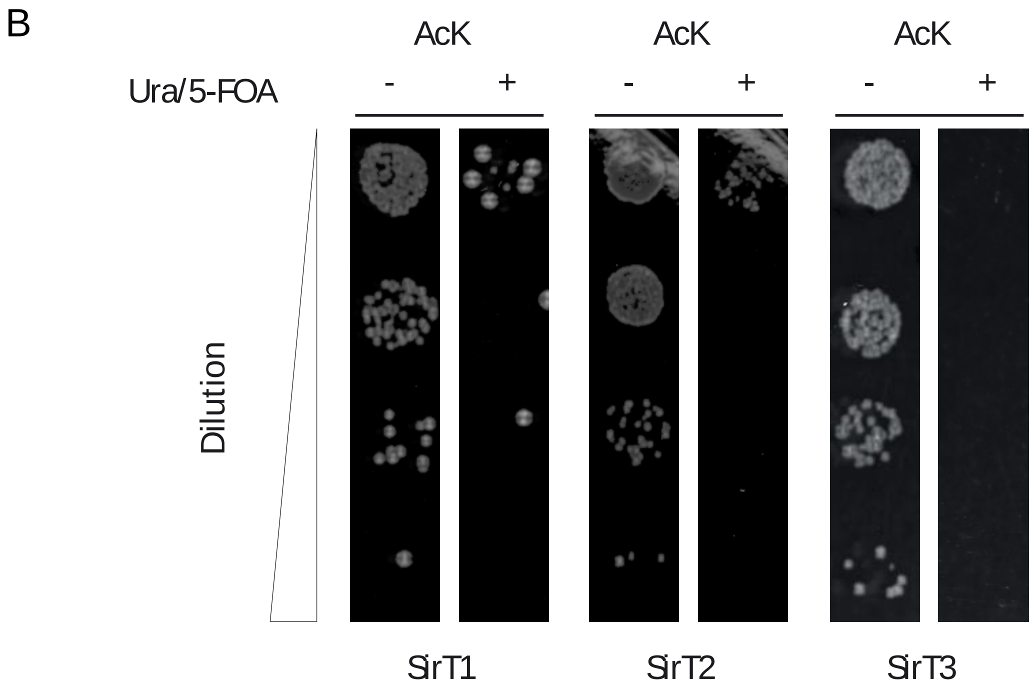

**Supplementary Figure 1: Selection of KDACs using Ura3 modified on K93**. **A**) DH10B DpyrF DcobB expressing CobB or HDAC8 and either Ura3 K93ac or Ura3 K93BocK were grown on minimal medium with or without uracil and 0.1% 5-FOA. **B**) DH10B DpyrF DcobB expressing one of three human Sirtuins and Ura3 K93ac were grown under conditions as in A.

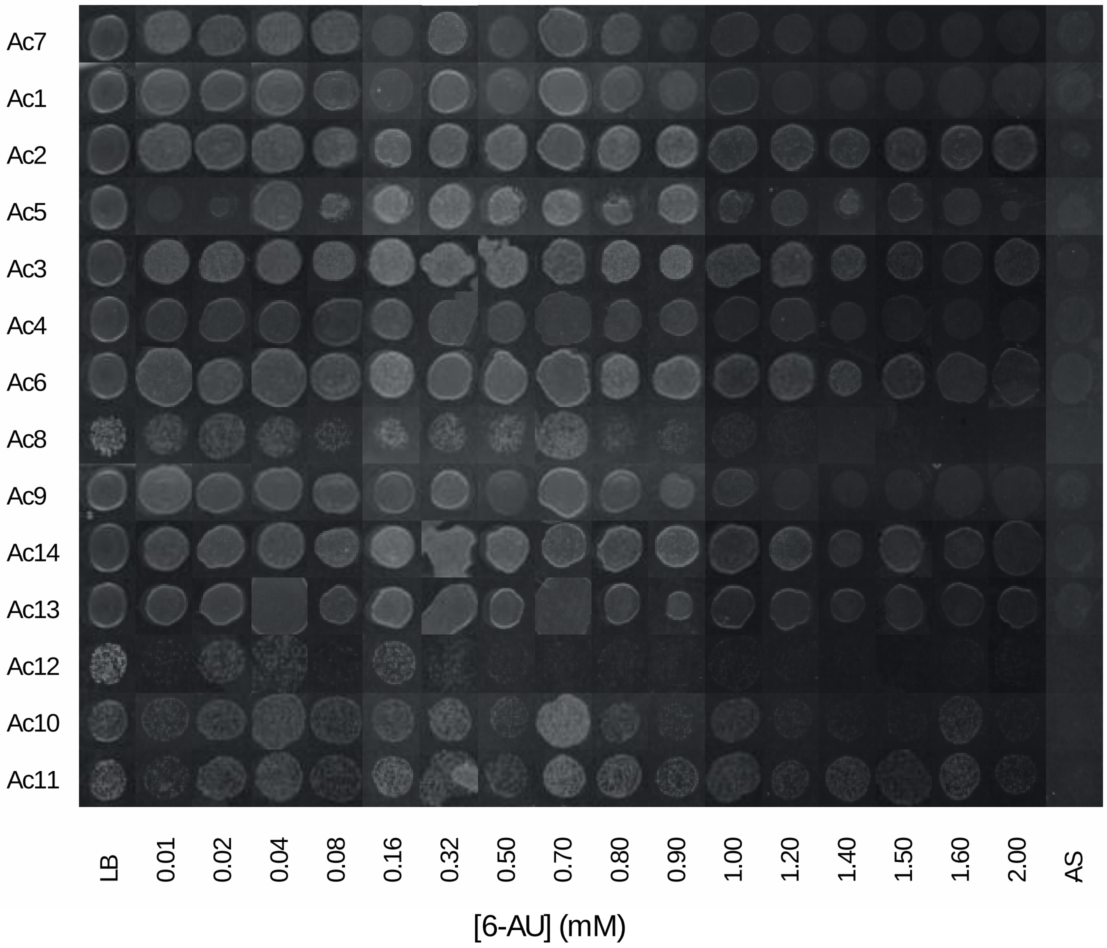

**Supplementary Figure 2: Growth of CobB variants in the presence of 6-AU.** E. coli expressing Ura3 K93ac as sole source of OMP decarboxylase and one of the CobB variants identified in the selection for acetyl-specificity were challenged to grow on LB agar plates in the presence of increasing concentrations of 6-AU.

A

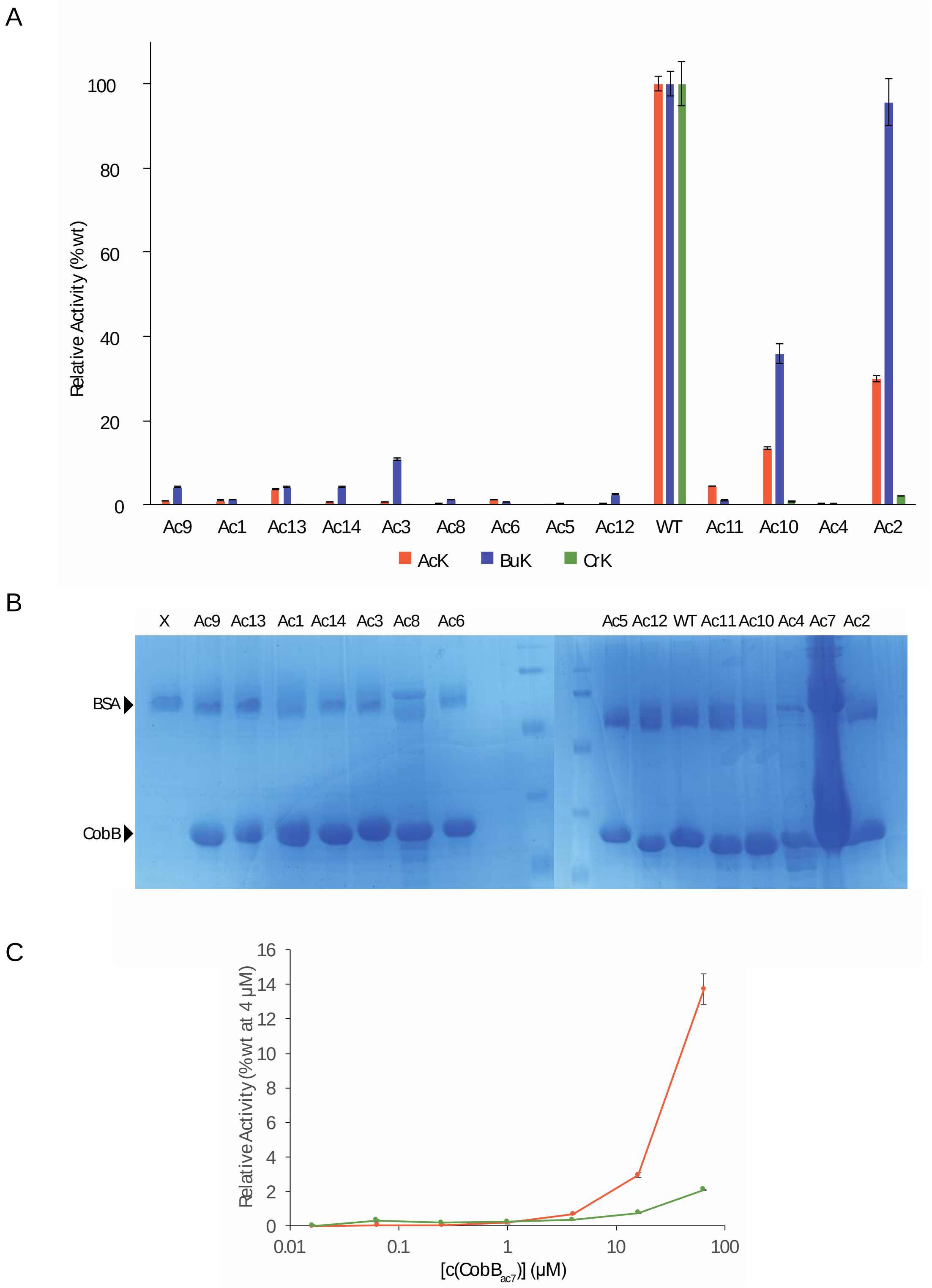

**Supplementary Figure 3: In vitro firefly luciferase assay to measure CobB variants activity.** **A**) Deacetylation, Debutyrylation and Decrotonylation activity of 14 CobB variants purified on Ni-beads tested on FLuc K529ac, K529bu and K529cr. **B**) The SDS-PAGE shows CobB proteins used in A. BSA was present during the deacylation reaction. **C**) Activity of CobBac7 towards FLuc K529ac and K529cr relative to CobB wild-type at 4 µM. CobBac7 was measured separately because of an error in the protein concentration measurement in experiment A.

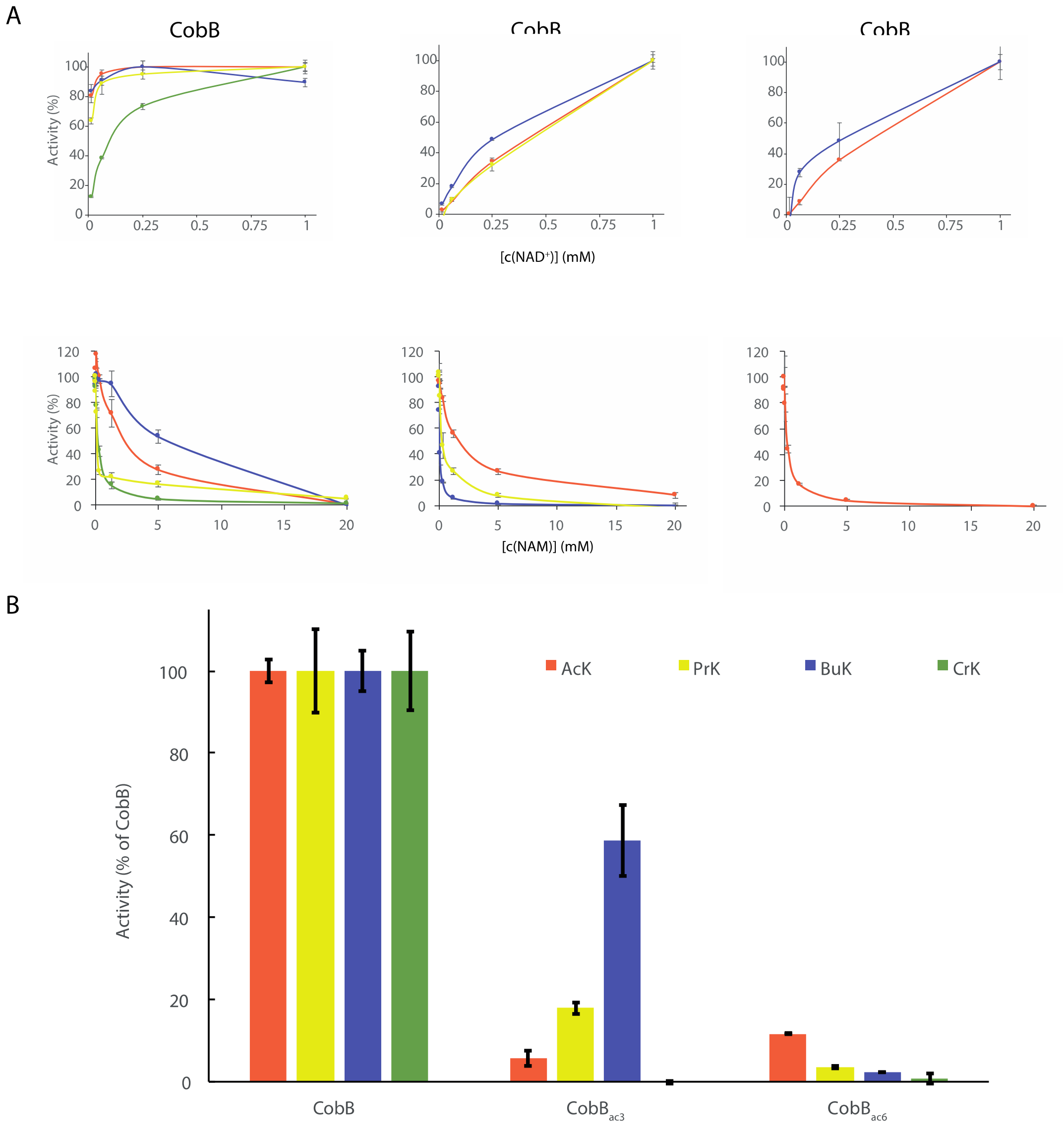

**Supplementary Figure 4: Catalytic rates of acetyl-selective CobB variants in dependence of NAD, NAM concentration and FLuc acylation. A**) FLuc with the indicated modification was incubated with CobB (64 nM), CobBac3 (2 µM) or CobBac6 (2 µM) and luminescence assayed in endpoint format. 2 mM NAD+ was present in the NAM titrations. **B**) CobB variants were assayed on purified FLuc carrying the indicated modification on K529. Experiments were performed in triplicates, error bars are standard deviation of the mean.

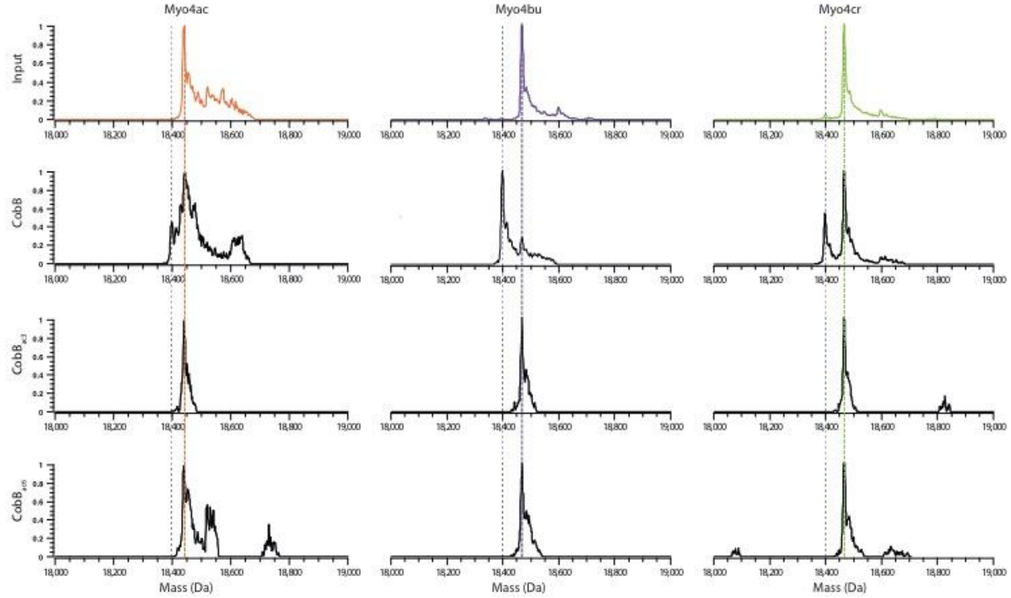

**Supplementary Figure 5: LC-ESI-MS analysis of Myoglobin proteins before and after incubation with CobB variants.** Dotted lines mark the expected mass of the free (grey) or modified Lysine (colored). No reaction was observed for $CobB_{ac3}$ (32 μM) and $CobB_{ac6}$ (16 μM) after 2 h incubation at 30°C by ESI-MS positive ion mode.

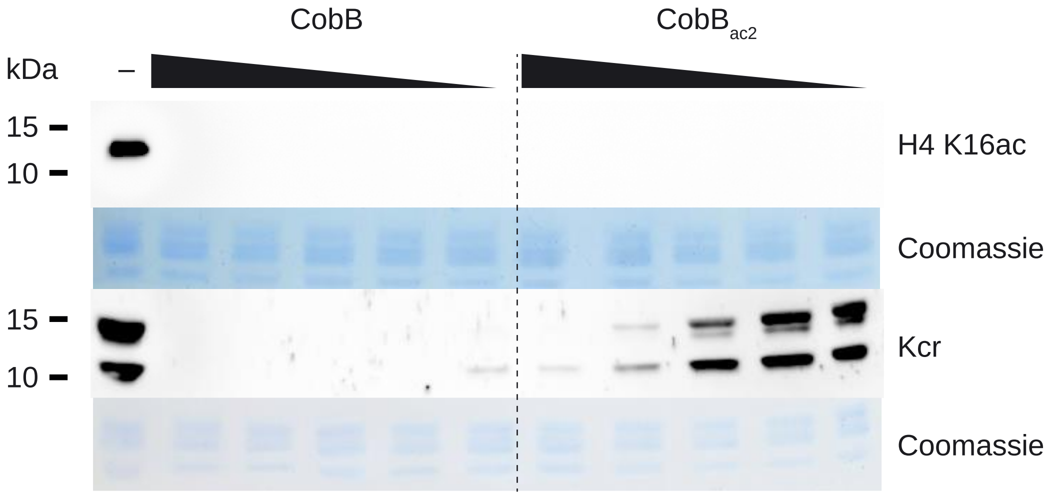

**Supplementary Figure 6: Deacylation of histones with CobB and variant CobBac2.** Purified human histones were incubated with CobB or CobBac2 (63 nM to 1 µM) for 2 h at 30°C in the presence of 2 mM NAD+. The modification state of the histones was analysed by Western blot using anti-H4 K16ac and anti-crotonyl-lysine antibodies**.**

A

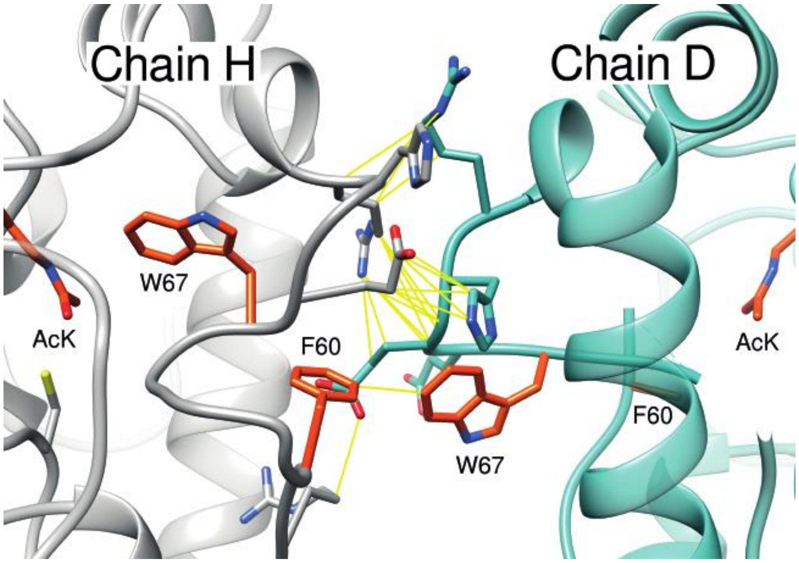

B

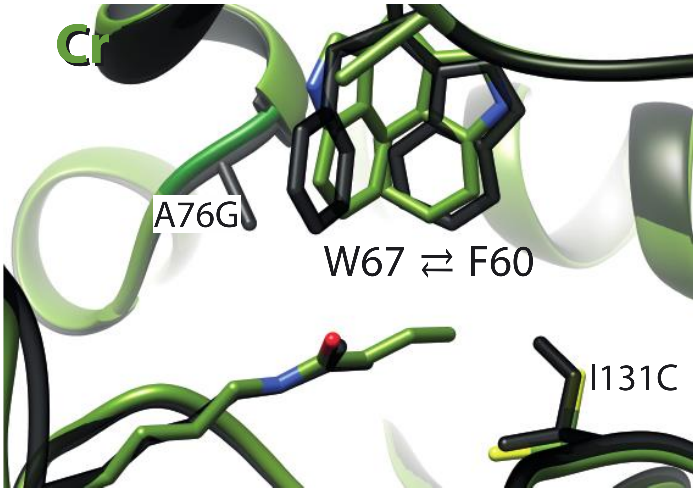

**Supplementary Figure 7: Omitted chains from overlay of CobB and CobB$_{ac2}$ A)** Crystal chains omitted from overlay of CobB$_{ac2}$ acetyl bound state (Figure 3C), due to contact (yellow lines) between the substrate binding loops of both chains. Acetyl-lysine, W60 and F60 of both chains (H, grey, D, opal) are highlighted in orange. **B**) Second molecule (Chain A) of CobB$_{ac2}$ in unit cell of H4 K16cr co-crystal. Chain A shows a mixture of conformations with W67 switching positions between the alternative and native state; F60 unresolved.

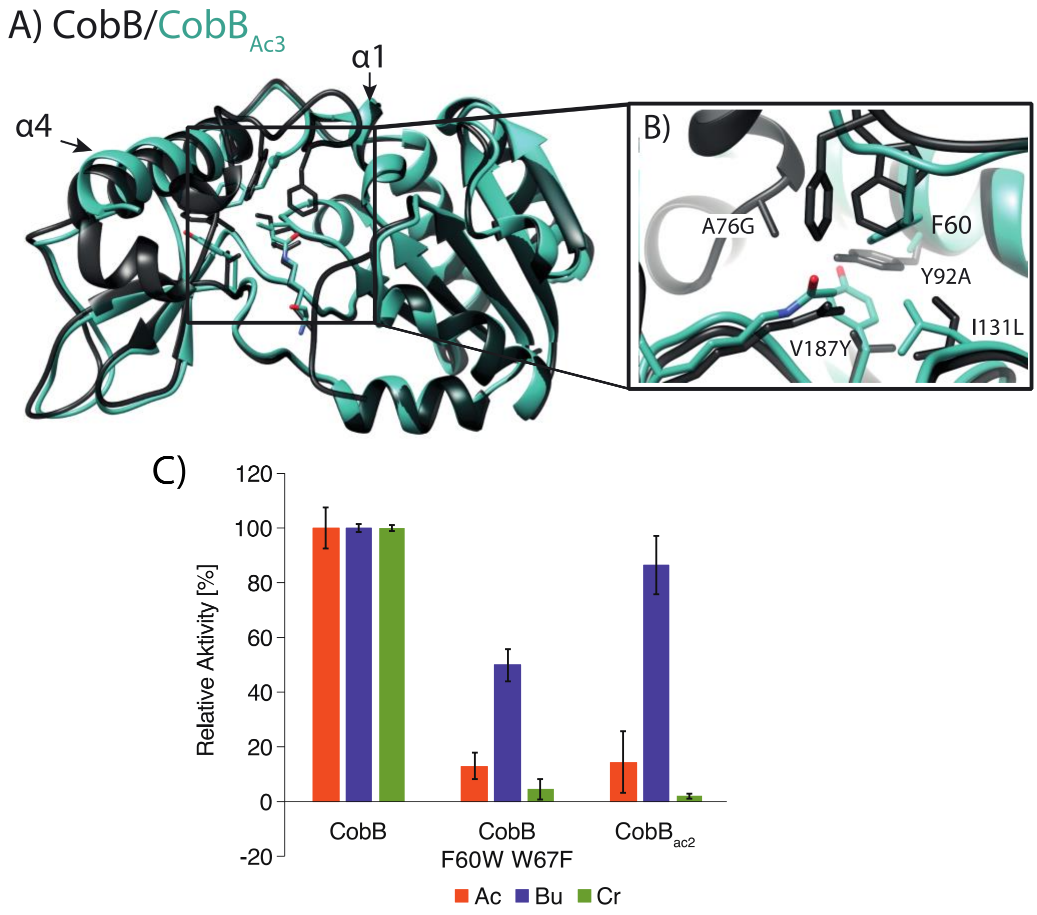

**Supplementary Figure 8: Structure of CobB$_{ac3}$ and CobB mimic mutants. A**) Overlay of H4 K16ac bound structures of CobB (black) and CobB$_{ac3}$ (A76G, Y92A, I131L, V187Y, opal). CobB$_{ac3}$ shows major distortions of cofactor binding loop and helix α2 and α3 following residue F60 until the beginning of helix α4. **B**) Close-up of the active site. The 4 mutations in CobB$_{ac3}$ allow F60 to bind the hydrophobic pocket previously held by W67. **C**) Selectivity of designed mutants (2 μM**)** towards different lysine modifications relative to CobB measured by Firefly luciferases KDAC assay for Acetyl (Ac) Buturyl (Bu) and Crotonyl-Lysine (Cr). Swapping F60 with W67 in CobB leads to acyl selectivity comparable to CobBac2. Error bars are standard deviations of the means of three biological replicates performed in triplicates.

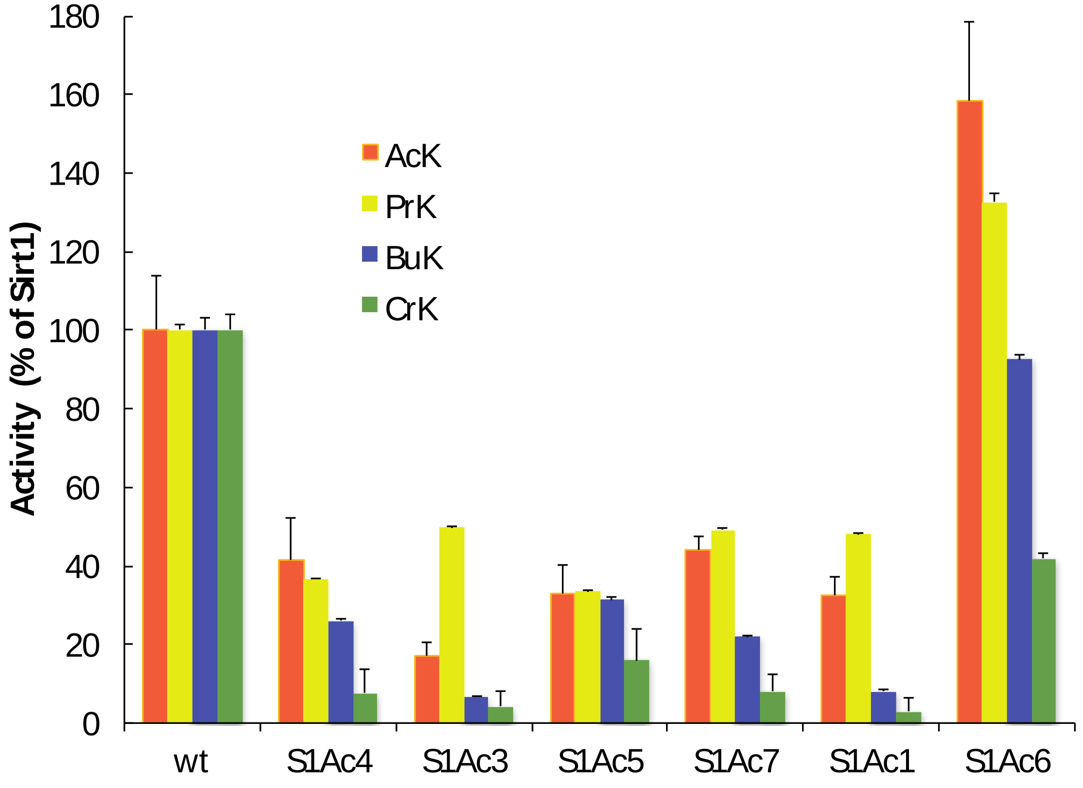

**Supplementary Figure 9: Acyl selectively of SirT1 variants.** SirT1 variants assayed on FLuc K529ac/pr/bu/cr. Error bars are standard deviations of the means of three replicates.

**Table S1**: CobB variants selected for AcK and against CrK cleavage.

| *Column1* | *A76* | *Y92* | *R95* | *I131* | *V187* | *Other* | *n* |
|---|---|---|---|---|---|---|---|
| $CobB_{ac1}$ | C | S | G | V | C | | *8* |
| $CobB_{ac2}$ | G | | | C | | V161A | *7* |
| $CobB_{ac3}$ | G | A | | L | Y | | *4* |
| $CobB_{ac4}$ | L | A | E | L | A | P36S, V45A, A203T | *4* |
| $CobB_{ac5}$ | L | K | W | L | I | | *5* |
| $CobB_{ac6}$ | L | S | S | L | W | | *4* |
| $CobB_{ac7}$ | S | G | K | | L | S206P | *18* |
| $CobB_{ac8}$ | S | I | A | | V | | *3* |
| $CobB_{ac9}$ | L | G | M | M | P | | *2* |
| $CobB_{ac10}$ | L | A | | L | P | | *1* |
| $CobB_{ac11}$ | L | | V | M | | | *1* |
| $CobB_{ac12}$ | L | M | T | V | Y | E103G, V161A | *1* |
| $CobB_{ac13}$ | L | W | N | W | Q | E79G, K192R, M202I | *1* |
| $CobB_{ac14}$ | V | V | N | W | Q | Q105R, D172G | *1* |
| *SUM* | | | | | | | *60* |

**Table S2**: Data collection and refinement statistics

| | **$CobB_{ac3}$-H4K16Ac** | **CobB-H4K16Ac** | **CobB-H4K16Bu** | **CobB-H4K16Cr** | **$CobB_{ac2}$-H4K16Bu** |
|---|---|---|---|---|---|
| **Space group** | C121 (5) | C2221(20) | P41212 (92) | P41212 (92) | P212121 (19) |
| **Cell parameters (Å)** | 90.25, 79.10, 39.90 | 131.10, 131.26, 58.54 | 92.73 92.73 58.62 | 93.96, 93.96, 58.54 | 60.49, 92.44, 94.45 |
| **(°)** | 90, 110.77, 90 | 90, 90, 90 | 90, 90, 90 | 90, 90, 90 | 90, 90, 90 |
| **Wavelength (Å)** | 0.978 | 1 | 1 | 1 | 1 |
| **Resolution (Å)** | 50-1.6 (1.66-1.60) | 50-1.6 (1.66-1.60) | 50-1.35 (1.38-1.35) | 50-2.3 (2.38-2.3) | 50-1.95 (2.02-1.95) |
| **Unique reflections** | 34254 (2536) | 66812 (4841) | 57070 (4152) | 12129 (881) | 39338 (2868) |
| **Multiplicity** | 6.5 (6.9) | 13.0 (13.2) | 25.2 (25.6) | 25.6 (25.7) | 13.1 (13.2) |
| **CC 1/2** | 99.8 (86.3) | 99.9 (64.4) | 97.8 (77.5) | 99.9 (81.6) | 99.9 (75.3) |
| **Completeness (%)** | 98.9 (99) | 100 (99.5) | 100 (99.9) | 100 (100) | 100 (100) |
| **$I/\sigma_I$** | 17.6 (3.1) | 18.8 (1.6) | 19.0 (1.6) | 23.9 (2.5) | 18.0 (1.7) |
| **$R_{merge}$ (%)** | 10.1 (70.4) | 6.7 (158.1) | 9.0 (261) | 9.8 (164.1) | 7.9 (163.7) |
| **$R_{work}/R_{free}$ (%)** | 17.9 / 20.5 | 17.2/18.7 | 16.9/18.8 | 21.1/25.0 | 19.7/21.5 |
| **Wilson B ($Å^2$)** | 14.7 | 27.8 | 18.4 | 53 | 39.3 |
| **Mean isotropic B factor ($Å^2$)** | 24.4 | 36.27 | 25.5 | 59.1 | 47.9 |
| **Bond length r.m.s.d. (Å)** | 0.010 | 0.008 | 0.012 | 0.003 | 0.004 |
| **Bond angle r.m.s.d. (°)** | 1.290 | 1.16 | 1.42 | 0.76 | 0.861 |
| **Ramachandran favoured (outliers)** | 100 (0) | 100 (0) | 98.0 (0) | 99 (0) | 98.6 (0) |
| **Number of atoms** | 3773 | 7800 | 4129 | 1931 | 4020 |
| **No H** | 1771 | 3707 | 2224 | 0 | 0 |
| **Water** | 220 | 294 | 287 | 27 | 187 |
| **Ligand (CIF)** | Acetyl-Lysine (aly) | Acetyl-Lysine (aly) | Butyryl-Lysine (btk) | Crotonyl-Lysine (kcr) | Butyryl-Lysine (btk) |
| **Pdb accession code** | 6RXS | 6RXJ | 6RXK | 6RXL | 6RXO |

Highest resolution shell in parentheses

**Table S2**: cont.

| | **$CobB_{ac2}$-H4K16Ac** | **$CobB_{ac2}$-H4K16Cr** | **$CobB_{ac2}$-H4K16Cr-Int3-36h** | **$CobB_{ac2}$-H4K16Cr-Int3-16h** |
|---|---|---|---|---|
| **Space group** | P212121 (19) | P212121 (19) | P1211 (4) | C121 (5) |
| **Cell parameters (Å)** | 92.30, 95.38, 168.83 | 57.56, 91.89, 95.90 | 61.38, 94.07, 94.06 | 132.9, 132.95, 57.99 |
| **(°)** | 90, 90, 90 | 90, 90, 90 | 90, 90.03, 90 | 90, 90.04, 90 |
| **Wavelength (Å)** | 1 | 1 | 1 | 1 |
| **Resolution (Å)** | 50-1.92 (1.99-1.92) | 50-1.8 (1.86-1.80) | 50-1.7 (1.76-1.70) | 50-2.0 (2.07-2.00) |
| **Unique reflections** | 114176 (8343) | 47854 (3481) | 117292 (8604) | 62989 (5013) |
| **Multiplicity** | 12.9 (12.4) | 12.6 (12.4) | 6.7 (6.9) | 6.9 (7.0) |
| **CC 1/2** | 99.8 (67.9) | 99.9 (83.5) | 100 (63.9) | 100 (68.5) |
| **Completeness (%)** | 100 (100) | 100 (99.9) | 100 (99.9) | 99.9(100) |
| **$I/\sigma_I$** | 13.9 (2.1) | 15.3 (2.1) | 20.1 (1.3) | 19.6 (1.8) |
| **$R_{merge}$ (%)** | 12.0 (133.0) | 8.5 (134.4) | 4.7 (142.5) | 4.3 (109.0) |
| **$R_{work}/R_{free}$ (%)** | 20.4/24.3 | 19.2/23.4 | 19.9/22.8 | 22.3 /25.9 |
| **Wilson B ($Å^2$)** | 32.3 | 30.7 | 32.9 | 48 |
| **Mean isotropic B factor ($Å^2$)** | 41.6 | 38.9 | 44.1 | 59.6 |
| **Bond length r.m.s.d. (Å)** | 0.010 | 0.007 | 0.007 | 0.004 |
| **Bond angle r.m.s.d. (°)** | 1.300 | 1.120 | 1.380 | 1.04 |
| **Ramachandran favoured (outliers)** | 99 (0.2) | 99.0 (0.2) | 97.0 (0.1) | 97.0 (0) |
| **Number of atoms** | 12180 | 4166 | 8316 | 8133 |
| **No H** | 0 | 0 | 0 | 0 |
| **Water** | 879 | 335 | 582 | 235 |
| **Ligand (CIF)** | Acetyl-Lysine (aly) | Crotonyl-Lysine (kcr) | Cr-Int.3 (2I3) | Cr-Int.3 (2I3) |
| **Pdb accession code** | 6RXM | 6RXP | 6RXQ | 6RXR |

Highest resolution shell in parentheses

**References**

1. Neumann H, Peak-Chew SY, & Chin JW (2008) Genetically encoding N(epsilon)-acetyllysine in recombinant proteins. *Nat Chem Biol* 4(4):232-234.
2. Neumann H, *et al.* (2009) A Method for Genetically Installing Site-Specific Acetylation in Recombinant Histones Defines the Effects of H3 K56 Acetylation. *Molecular Cell* 36(1):153-163.
3. Neumann H, *et al.* (2009) A method for genetically installing site-specific acetylation in recombinant histones defines the effects of H3 K56 acetylation. *Mol Cell* 36(1):153-163.
4. Bryson DI, *et al.* (2017) Continuous directed evolution of aminoacyl-tRNA synthetases. *Nature chemical biology* 13(12):1253-1260.
5. Heitmuller S, Neumann-Staubitz P, Herrfurth C, Feussner I, & Neumann H (2018) Cellular substrate limitations of lysine acetylation turnover by sirtuins investigated with engineered futile cycle enzymes. *Metab Eng* 47:453-462.
6. Knyphausen P, *et al.* (2016) Insights into Lysine Deacetylation of Natively Folded Substrate Proteins by Sirtuins. *J Biol Chem* 291(28):14677-14694.
7. Jiang W, Bikard D, Cox D, Zhang F, & Marraffini LA (2013) RNA-guided editing of bacterial genomes using CRISPR-Cas systems. *Nature biotechnology* 31:233.
8. Gabriella S, *et al.* (2006) The Mechanism of Plasmid Curing in Bacteria. *Current drug targets* 7(7):823-841.
9. Shechter D, Dormann HL, Allis CD, & Hake SB (2007) Extraction, purification and analysis of histones. *Nature Protocols* 2:1445.
10. Zhao K, Chai X, & Marmorstein R (2004) Structure and substrate binding properties of cobB, a Sir2 homolog protein deacetylase from Escherichia coli. *J Mol Biol* 337(3):731-741.
11. Kabsch W (2010) Xds. *Acta Crystallogr D Biol Crystallogr* 66(Pt 2):125-132.
12. McCoy AJ, *et al.* (2007) Phaser crystallographic software. *J Appl Crystallogr* 40(Pt 4):658-674.
13. Murshudov GN, Vagin AA, & Dodson EJ (1997) Refinement of macromolecular structures by the maximum-likelihood method. *Acta Crystallogr D Biol Crystallogr* 53(Pt 3):240-255.
14. Adams PD, *et al.* (2010) PHENIX: a comprehensive Python-based system for macromolecular structure solution. *Acta Crystallogr D Biol Crystallogr* 66(Pt 2):213-221.
15. Lebedev AA, *et al.* (2012) JLigand: a graphical tool for the CCP4 template-restraint library. *Acta Crystallogr D Biol Crystallogr* 68(Pt 4):431-440.
16. van Aalten DM, *et al.* (1996) PRODRG, a program for generating molecular topologies and unique molecular descriptors from coordinates of small molecules. *J Comput Aided Mol Des* 10(3):255-262.
17. Emsley P, Lohkamp B, Scott WG, & Cowtan K (2010) Features and development of Coot. *Acta Crystallogr D Biol Crystallogr* 66(Pt 4):486-501.
